# Temporal Gating by Chandelier Cells Encodes Signed Prediction Errors

**DOI:** 10.64898/2026.06.26.734797

**Authors:** Przemyslaw Jarzebowski, Daniel Bendor

## Abstract

The brain refines its predictions of the world by updating its internal model whenever sensory input differs from expectation. The sign of this prediction error matters: an unexpected event signals that the model under-predicted (positive error), while a predicted event that fails to occur indicates that the model over-predicted (negative error), and the two should drive opposite synaptic changes. How cortical circuits represent error sign in spiking activity, and how that representation translates into synaptic learning, remain unresolved. We propose the Signed Error by Timing Asymmetry (SETA) model, in which the sign of a prediction error is encoded by *when* layer 2/3 neurons fire relative to a brief plasticity window in their layer 5 targets. Chandelier cells, an inhibitory cell type recruited by the prediction, impose a temporal clamp on layer 2/3 output: positive errors escape the clamp and arrive within the synaptic potentiation window, while negative errors are released only after the clamp decays and arrive later, during the synaptic depression window. The same circuit, therefore, biases downstream synapses toward either potentiation or depression depending on the prediction-error sign. We demonstrate this signed-error computation in a reduced two-compartment model, test SETA-specific predictions using in vivo recordings from mouse visual cortex, and examine how E/I imbalance leads to pathological consequences in predictive coding.

## INTRODUCTION

Predictive coding has emerged as the leading theoretical framework for cortical computation, proposing that the brain continuously generates predictions about incoming sensory input and refines its internal model using prediction errors - the difference between prediction and sensation (Rao & Ballard 1999; Friston 2005; Keller et al. 2012). This framework provides a principled account of how the brain learns efficiently from temporally structured input, unifying cortical phenomena ranging from the suppression of responses to repeated stimuli to the amplification of signals that violate learned regularities. The framework’s central computational claim is that prediction errors must drive plasticity in the internal model, and that the influence of any sensory signal on this update is scaled by its surprise value (Friston 2010). Yet how cortical circuits represent the sign of a prediction error in spiking activity, and how that sign determines the direction of plasticity at downstream synapses, remains unresolved.

The laminar organisation of the neocortex offers a natural scaffold. Bottom-up sensory drive arrives via thalamus → Layer 4 → Layer 2/3, while top-down signals converge on apical dendrites via Layer 1 (Bastos et al. 2012, 2020). Layer 2/3 has been proposed as the primary site of prediction error computation, with errors hierarchically forwarded to update higher-order representations. Errors can be positive (sensation > prediction) or negative (prediction > sensation), with surprise and omission as their extremes. Single-population rate codes face a representational asymmetry: positive errors can be expressed as firing rate increases above baseline, whereas negative errors require sustained firing rate decreases that are difficult to maintain given the low baseline firing rate of L2/3 neurons (Sakata and Harris 2009). Keller and Mrsic-Flogel (2018) addressed this with a dual-population scheme of distinct positive- and negative-error neurons, formalised by Hertäg and Clopath (2022) and supported by cell-type-specific imaging (Attinger et al. 2017; Jordan & Keller 2020; O’Toole et al. 2023). However, rate-coding accounts do not specify how prediction errors drive the long-term plasticity necessary for learning and maintaining predictive models. Cortical plasticity rules for bidirectional synaptic change rely on millisecond timing of spikes relative to dendritic calcium events, rather than solely on time-averaged firing rates (Sjöström and Häusser 2006; Letzkus et al. 2006; Graupner and Brunel 2012).

### Translation of prediction-error sign to synaptic plasticity direction

We propose that Layer 5 extratelencephalic (ET) pyramidal neurons are a site translating prediction errors into directed synaptic change. Their dendrites segregate inputs into distinct compartments: basal dendrites integrate sensory inputs (Constantinople and Bruno 2013) and local recurrent drive from L5 intratelencephalic (IT) neurons (Kim et al. 2015), while the apical tuft in Layer 1 receives signals from higher-order cortical areas (Petreanu et al. 2009; Harris and Shepherd 2015). We propose that the recurrent L5 IT network is an ideal candidate for representing internal predictions. Its nodes are interconnected across the cortex (Heindorf and Keller 2024) and provide a local relay of a prediction signal in the model. Either prediction or sensory inputs arriving at the basal compartment can generate an action potential in L5 ET neurons that leads to a backpropagating action potential (bAP). If the generated bAP is near-coincident with apical tuft depolarisation, a backpropagation-activated calcium (BAC) firing is facilitated, leading to a somatic burst (Fig. 1A; Larkum et al. 1999; Larkum 2013). BAC firing links these two compartments, strongly potentiating recently active apical synapses; non-coincident activation favours their depression or no change (Sjöström & Häusser 2006; Letzkus et al. 2006). However, learning of this association must be gated by sensory evidence, not just an internal prediction in the absence of sensory input. We propose that learning is controlled by the oblique dendrites, situated between soma and tuft, as they can facilitate BAC initiation during the acquisition of new associations. This is because when tuft synapses are weak, oblique input acts synergistically with the bAP, boosting the forward spread of distally initiated potentials and supporting BAC firing (Larkum et al. 2001; Schaefer et al. 2003). Oblique input arriving before the bAP has decayed biases tuft plasticity toward LTP; late input raises intracellular calcium toward the LTD range without triggering BAC firing (Graupner and Brunel 2012; Paulsen and Sejnowski 2000). This bAP-initiated plasticity window due to an increased probability of BAC firing (Larkum et al. 1999), provides a countdown for potential facilitation by the oblique input. Because L2/3 projects to the oblique dendrites of L5 ET neurons (Petreanu et al. 2009, Zhang et al. 2026), it is anatomically positioned to gate BAC firing, and hence the plasticity direction. We postulate that a L2/3 prediction error signal delivered to the oblique dendrites at the correct time can facilitate LTP of the tuft for underpredictions and its LTD for overpredictions, translating L2/3 prediction errors into directed synaptic change.

**Figure 1:**
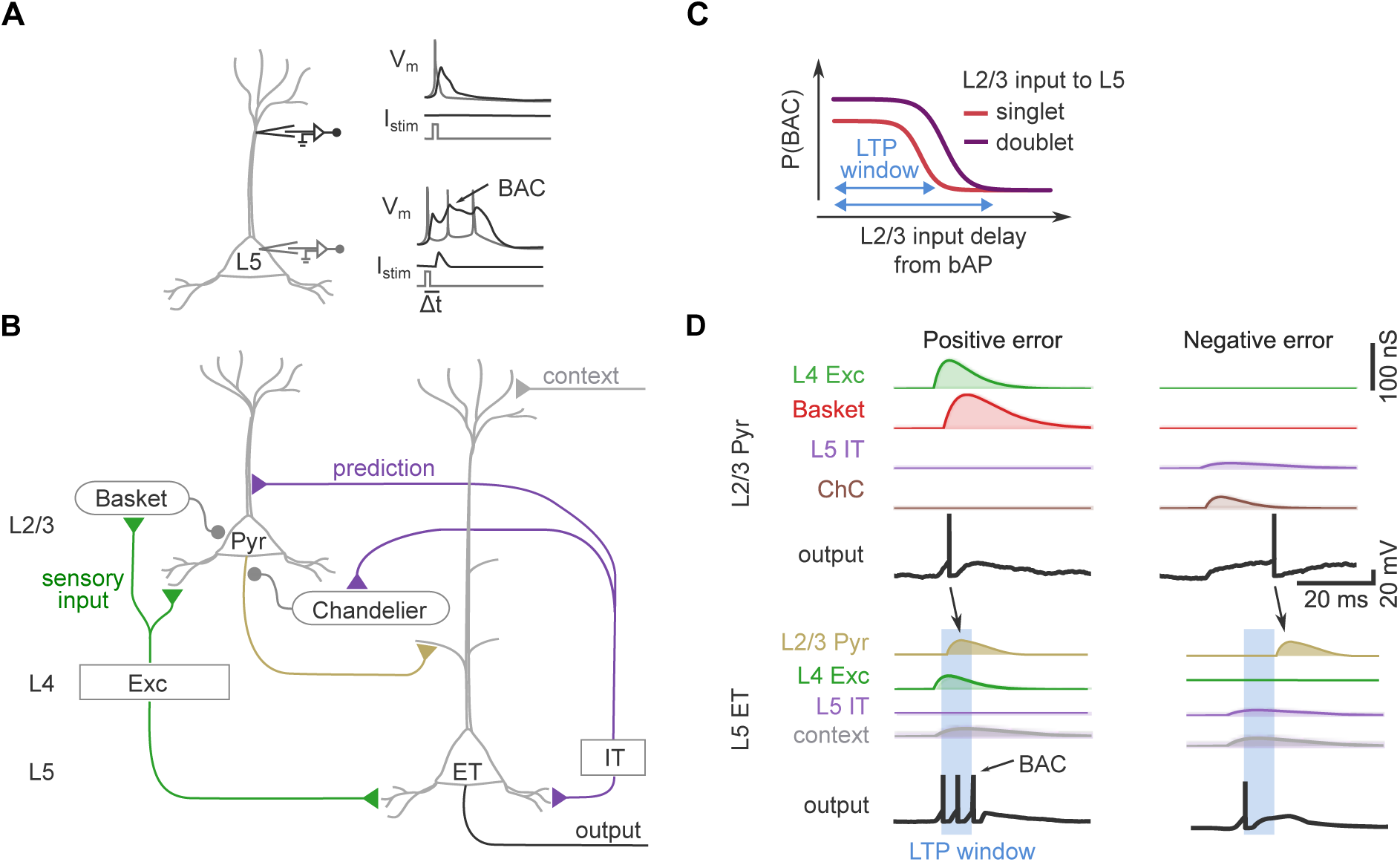
Cortical circuit and mechanism for updating synaptic weights based on signed prediction error. (A) Coincident somatic and distal depolarisation of L5 pyramidal neuron facilitates BAC firing. Current injected at the distal location (black pipette, left panel) when coupled with injection at the soma (grey pipette, left panel) facilitates BAC generation and a somatic burst (bottom right panel), compared to somatic current injection alone (top right panel). Figure reproduced from Larkum et al. (1999). (B) Schematic of the cortical circuit that temporally gates L2/3 input onto L5 ET neuron. L2/3 pyramidal cell integrates sensory and prediction signals. Its spike timing is gated by two inhibitory cell types: Basket cells targeting the soma, and Chandelier cell targeting axon initial segment. The L2/3 spike timing gates whether L5 ET neuron will strengthen or weaken the association between predictive signal in its basal dendrites and the L1 contextual input arriving at its apical tuft. (C) Schematic showing how the timing and strength of L2/3 inputs that follow bAP in the L5 ET neuron affect the likelihood of initiating BAC firing. (D) Simulation of two-compartment neuron model that temporally gates L2/3 spiking. When the sensory signal drives L4 Excitatory cell and Basket cell conductance in absence of predictive conductance (positive error), the L2/3 neuron generates a short-latency spike. This spike arrives after the L5 ET neuron fires, falling within window for BAC generation (LTP window). A positive prediction error results in LTP at two sites: (1) at the tuft synapses associating the contextual input with the predictive signal arriving directly at the L5 ET neuron, and (2) at the L2/3 oblique synapses, increasing gating strength. Conversely, when a prediction signal from L5 IT drives L2/3 and Chandelier cell (ChC) conductance, in the absence of sensory conductance (negative error), Chandelier cell temporally clamps the potential at the L2/3 axon initial segment below spike-threshold. When the clamp is released, L2/3 neuron fires a spike, delayed relative to the L5 IT input. This spike has a low likelihood of generating BAC in the L5 ET neuron. Consequently, this negative prediction error results in LTD of the tuft synapses, weakening the association in the L5 ET neuron between the predictive signal arriving from the L5 IT neuron and the coincident contextual input.

### Representation of prediction-error sign in L2/3 spike timing

We propose a model representing signed prediction error based on the precise timing of the L2/3 spiking relative to L5 ET spiking. Their spike timing is mediated by their shared inputs. Both L2/3 and L5 ET neurons receive inputs from L4 and L5 IT (Hage et al. 2022; Brown and Hestrin 2009; Petreanu et al. 2009, Zhang et al. 2026). These parallel pathways relay shared sensory (L4) and predictive (L5 IT) inputs (Fig. 1B), synchronizing the spiking of the L2/3 neuron to the L5 ET neuron spike that triggers the bAP-initiated plasticity window. The exact L2/3 spike time controls the direction of synaptic plasticity: on underprediction, the minimal delay of the L2/3 input arriving at the L5 ET oblique facilitates BAC firing, while on overprediction, a long enough delay causes the L2/3 spike to fall outside the LTP-biased portion of the plasticity window, biasing the synaptic change towards LTD. L4 and L5 IT neurons each recruit a distinct parvalbumin (PV) interneuron subtype (Fig. 1B; Helmstaedter et al. 2008; Xu and Callaway 2009; Seignette et al. 2024) that we propose is central to the spike-timing control that enables prediction error calculation. The first PV interneuron subtype, chandelier cells (ChCs), target the axon initial segment of L2/3 pyramidal neurons (Somogyi 1977) and exert a depolarising subthreshold clamp (Woodruff et al. 2009, 2011). Evidence suggests that ChCs are recruited by L5 IT neurons (Xu and Callaway 2009; Seignette et al. 2024) and, in our model, they can prevent L2/3 spike initiation. If their conductance decays while the soma remains depolarised from the concurrent L5 IT or other inputs, a spike, albeit delayed, can be released. We refer to this as a gated release spike, whose latency encodes the strength of the prediction relative to the sensory input. A negative error should produce a delay exceeding the initial LTP-biased portion of the bAP-initiated plasticity window in L5 ET neurons, favouring LTD instead. The second PV interneuron subtype, basket cells, are driven by L4 sensory input, hyperpolarising the L2/3 pyramidal neurons’ soma with a ∼2–3 ms feedforward delay (Helmstaedter et al. 2008; Pala and Petersen 2015). Given their short delay relative to L4 input, they do not impede the initial sensory spike in the model but prevent subsequent longer latency spikes. Thus, basket cells contribute to the model encoding positive error in L2/3 neurons: L4-driven spikes fall within the initial LTP window and are actively prevented from firing later when LTD in the L5 ET neuron would be favoured. Together, these two PV interneuron subtypes implement a single-population mechanism for encoding prediction error sign in spike latency, which we term the Signed Error by Timing Asymmetry (SETA) model. The model is parsimonious: a single population, a single readout (the probability of BAC firing), and PV interneuron-gated spike timing suffice to translate the sign of prediction error into the direction of cortical plasticity.

Here, we formalise SETA in a two-compartment simulation, implement precision weighting of the prediction signal, and examine how distinct causes of E/I imbalance produce qualitatively different computational failures that are relevant to autism and schizophrenia. Lastly, we test the model’s key predictions against extracellular spiking recordings from mouse V1 published by Allen Brain Observatory.

## RESULTS

### A two-compartment model captures the differential roles of basket cells and chandelier cells in controlling L2/3 spike timing

To investigate how L2/3 pyramidal neurons integrate prediction and sensory input to compute a signed prediction error, we developed a leaky integrate-and-fire (LIF) two-compartment model separating the somatic and axon initial segment (AIS) compartments. This architecture was chosen specifically to capture the differential consequences of somatic inhibition by basket cells versus axo-axonic inhibition by chandelier cells (ChCs), a distinction that is critical to the temporal gating mechanism at the core of the SETA model. In the somatic compartment, the model integrates three synaptic streams: fast AMPA-mediated bottom-up sensory drive from L4, local predictive input from L5 IT neurons, and fast feedforward inhibitory input from basket cells. The AIS compartment receives targeted axo-axonic inhibition from ChCs, modelled with a depolarising GABA_A_ reversal potential (E_ChC_ = −58 mV), close to but below spike threshold (Woodruff et al. 2009, 2011). Axial conductance couples the two compartments bidirectionally, allowing somatic depolarisation to accumulate near the AIS even while ChC conductance is active. ChC input creates what we term a temporal clamp: somatic depolarisation is transferred axially to the AIS where it is held below spike threshold by the ongoing shunting conductance. This behaves like a compressed spring: when the ChC conductance decays, the somatic drive pushes the AIS past spike threshold, producing a gated release spike whose timing is set by the ChC conductance kinetics rather than the temporal precision of the excitatory somatic drive. This transforms the question of whether the neuron fires into a question of when, making the timing of L2/3 output available as a continuous variable encoding the error sign.

#### Positive prediction error: fast sensory drive synchronises L2/3 spike with L5 bAP

When L4 sensory input arrives in the absence of L5 IT predictive input, the L2/3 neuron produces a singlet or doublet depending on the L4 input strength. L4 drive depolarises the soma rapidly; basket cells, also driven by L4 input, engage with a latency of 2.5 ms and provide hyperpolarising somatic inhibition that terminates the response (Helmstaedter et al. 2008). Because no ChC clamp is active, action potential initiation at the AIS is rapid and unimpeded. The resulting spike or burst occurs immediately following sensory input arrival, reaching the L5 ET oblique synapse in near coincidence with the bAP, synergistically increasing the likelihood of BAC firing (Larkum et al. 1999; Larkum 2013; Sjöström and Häusser 2006; Letzkus et al. 2006). For simplicity, we refer to the BAC-permissive window, in which L2/3 input increases the likelihood of LTP at L5 ET tuft synapses, as the LTP window. As L2/3 spike latency relative to the bAP increases, P(BAC) decreases, shifting the plasticity outcome continuously from LTP through no change toward LTD. In vitro, the ability of distal L5 inputs to facilitate BAC returns to baseline within approximately 15 ms post-bAP (Larkum et al. 1999). While the exact boundary between the LTP and LTD windows depends on the strength of multiple inputs; what matters for the model is that sensory-driven spikes fall within the LTP window and prediction-driven release spikes fall outside it. The L2/3 input arriving within the LTP window encodes a positive prediction error (Fig. 1D). A doublet of L2/3 spikes provides a stronger and more sustained oblique input than a singlet, enhancing depolarisation of the decaying bAP or triggering an additional L5 somatic spike, further increasing P(BAC) (Fig. 1C).

#### Negative prediction error: the axoaxonic gated release spike following L5 bAP

By contrast, when L5 IT predictive input arrives to the L2/3 pyramidal cell but the sensory input is absent or weaker than the prediction, the L2/3 cell produces a gated release singlet. L5 drive depolarises the soma and simultaneously recruits ChC (Xu and Callaway 2009; Seignette et al. 2024), clamping the AIS near threshold. In the absence of additional sensory drive from L4, the AIS cannot escape this clamp until ChC conductance sufficiently decays, at which point the accumulated depolarisation is released as a singlet if the soma remains sufficiently depolarised (Fig. 1D). In such case, the L2/3 input arrives to the L5 ET 15–20 ms delayed after the L5 neuron’s bAP could have been initiated by the same L5 IT predictive input that initiated the L2/3 spike. The delayed L2/3 input relative to the L5 ET bAP biases the plasticity towards LTD (Fig. 1C).

#### The circuit encodes a continuous error spectrum

We next simulated how a graded sensory input interacts with a ChC clamp of fixed strength. When L4 input was weaker than the predictive input, the ChC clamp successfully vetoed the L2/3 spike, and basket cell hyperpolarisation prevented the gated release spike (Fig. 2A), resulting in no spiking. When the L4 input was sufficiently strong and arrived during the active ChC clamp window, it overrode the clamp and drove a spike, a breakthrough event, albeit with a higher threshold for producing a doublet relative to a completely unpredicted positive error (Fig. 2B). The model was robust to exact time difference between prediction and sensory inputs and delaying either input by 5 ms resulted in the same behaviour (Fig. 2C). The ratio of sensory drive to predictive clamp also determined the latency of the breakthrough spike. A strong sensory drive (relative to the prediction) produced a near-coincident L2/3 spike with the sensory-driven bAP, encoding a positive error. As sensation weakened relative to prediction, breakthrough latency lengthened: the resulting L2/3 spike arrived progressively later relative to the bAP (Fig. 2D), sliding the expected plasticity outcome from potentiation towards depression along a continuous gradient rather than switching abruptly between them.

**Figure 2:**
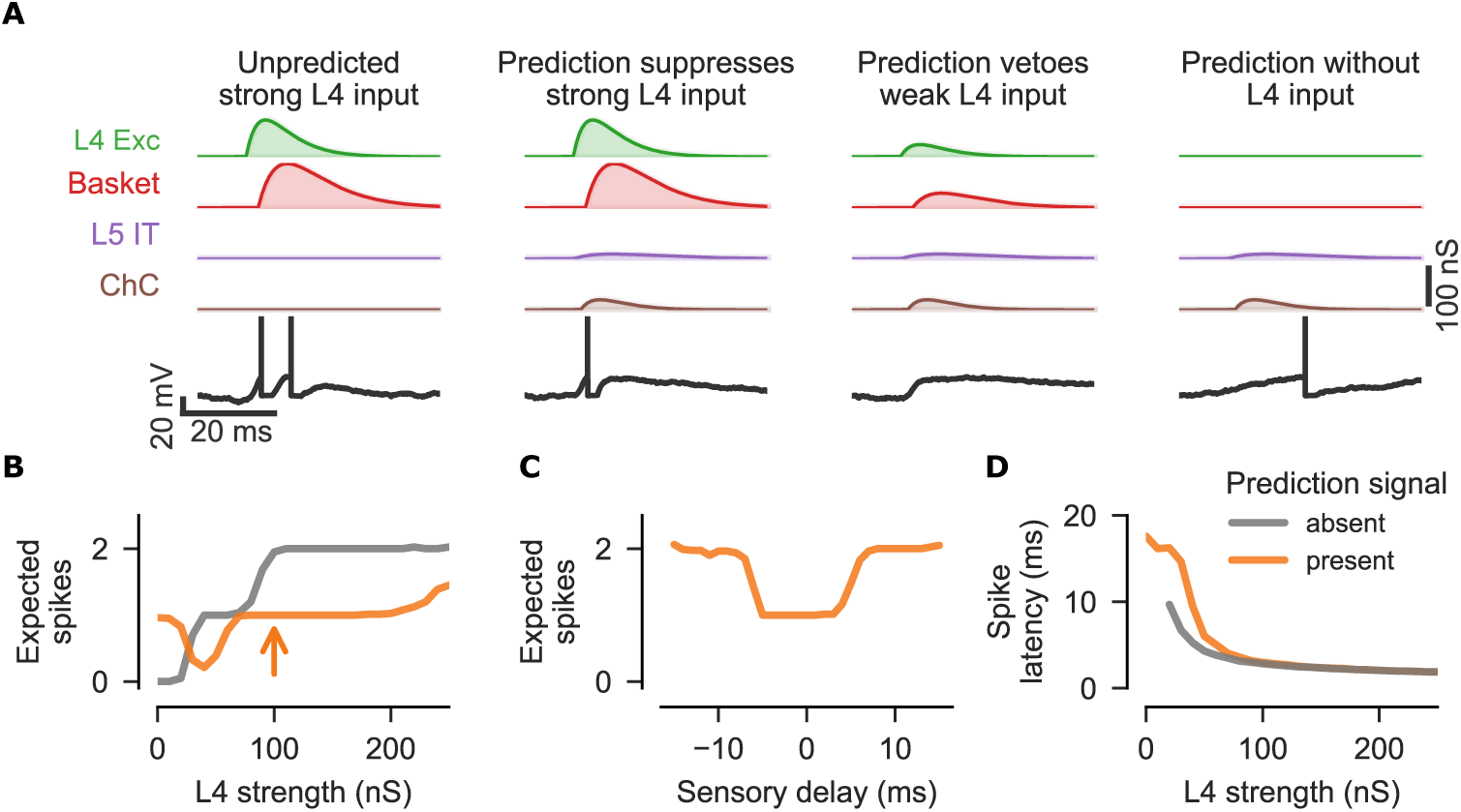
Interaction between prediction and sensory input. (A) Simulation of two-compartment neuron model of L2/3 spiking demonstrating how spike count and latency change with the relative conductances driven by sensory and predictive signals.(B) Expected L2/3 spike count as a function of L4 Excitatory cell conductance. The basket cell conductance was scaled proportionally. The plot compares the expected spike count when the sensory signal interacts with prediction signal of fixed strength (L5 IT conductance of 12 nS, ChC conductance of 25 nS) to a baseline without prediction conductances.(C) Expected spike count as a function of sensory input delay from predictive L5 IT conductance onset. The arrow in panel B marks the L4 Excitatory cell conductance in the simulation.(D) Latency to the first L2/3 spike from sensory input as a function of L4 Excitatory cell conductance for simulation shown in panel B.

The continuous gradient of spike latencies reframes what may constitute a “perfect prediction.” Previous predictive coding models have proposed that perfect prediction (sensory input strength = prediction input strength) corresponds to neuronal silence (Rao & Ballard 1999; Friston 2005, Keller & Mrsic-Flogel 2018). Within SETA, silence at L2/3 has more nuanced consequences: oblique synapses receive no input and therefore undergo no plasticity, but tuft synapses still experience the bAP-evoked calcium transient; in the absence of a BAC, this is more likely to drive LTD than LTP (Graupner and Brunel 2012). Silence is therefore not a plasticity-neutral state - it can weaken the L5 ET neuron’s apical tuft synapses even when predictions are accurate. A more meaningful definition of perfect prediction is a state in which the expected plasticity at recently active L5 ET tuft synapses is balanced over repeated trials, with P(LTP) ≈ P(LTD), so that the overall expected change is close to zero. Oblique synapses contributing to those events follow the same balance criterion. Because P(BAC) decreases with the latency of L2/3 input, a delayed L2/3 singlet can produce this balanced state across a range of conditions. Additionally, a decrease in spike reliability (expected spike count between 0 and 1 spike; Fig. 2B), may also contribute to this balance. Our simulation suggests that the prediction/sensation ratio determines the position along this gradient, with weak sensory relative to prediction inputs resulting in a longer latency L2/3 spike and hence shifting the outcome toward LTD and stronger sensory inputs toward LTP (Fig. 2D).

### Precision weighting of prediction error

The sign of the prediction error is necessary but not sufficient to drive appropriate learning. While the magnitude of the positive error scales with sensory drive, the learning rate controls how effectively positive and negative errors update downstream synaptic weights, and should be modulated by internal confidence and behavioural state, a requirement formalised as precision weighting (Friston 2010; Feldman and Friston 2010). Previous models have implemented precision weighting by scaling the synaptic gain of dedicated L2/3 prediction error units (Spratling 2008; Keller & Mrsic-Flogel 2018). SETA achieves the same effect by scaling of sensory and predictive gain at source. A change in the feedforward gain of L4 and L5 IT proportionally scales their downstream PV interneurons (basket and chandelier, respectively). A balanced increase of excitatory and inhibitory inputs increases the magnitude of the prediction error while maintaining its sign. Our model is agnostic to which mechanism sets the gain; we require only that excitation and inhibition co-scale. Arousal- and locomotion-related Vip activity is positioned to drive this modulation by disinhibiting the gain-setting Somatostatin-expressing (Sst) cell population, though the relationship is more complex than a single disinhibitory control point (Pi et al. 2013; Pfeffer et al. 2013; Polack et al. 2013; Bugeon et al. 2022). Following previous precision-weighting models (Spratling 2008; Keller & Mrsic-Flogel 2018), our simulation co-varied sensory and predictive gain, scaling the precision of positive and negative errors together. In principle the two streams could be controlled independently.

We simulated a graded somatic gain of L4 and L5 IT neurons, which proportionally scaled excitatory and inhibitory L2/3 inputs (see Methods). At low gain, both error signals were effectively suppressed: strong sensory inputs rarely drove bursts, and stimulus omission frequently ended in silence rather than a delayed gated release spike (Fig. 3A). As L4 and L5 IT gain increased the sensory threshold for bursts decreased (favouring LTP; Fig. 3A,B), and the probability of a gated release spike on omission trials increased (favouring LTD; Fig. 3B). Critically, increased gain did not substantially alter the delay of the gated release spike when it occurred (Fig. 3C). Therefore, the spike timing preserved sign of the error; gain modulated only the firing rate and reliability of the error signal.

**Figure 3:**
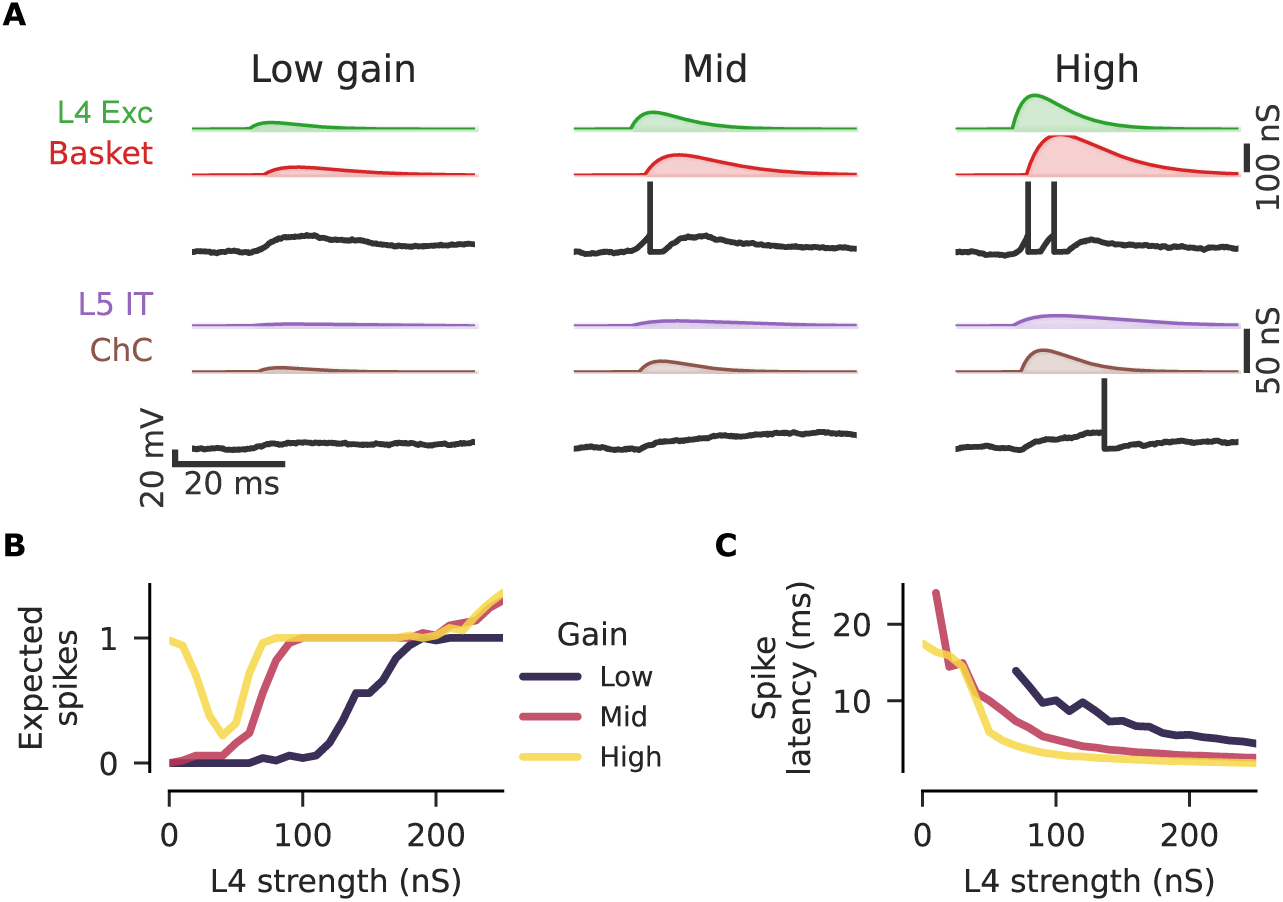
Precision weighting of prediction error. (A) Simulation of L2/3 firing across different L4 and L5 IT somatic gain levels. The simulation was run for two scenarios: (1) sensory conductances only (top panel), (2) predictive conductances only (bottom panel). Somatic gain was modelled by equal scaling of excitatory and inhibitory inputs set to 0.2 for Low, 0.5 for Mid, and 1.0 for High gain.(B) Expected L2/3 spike count as a function of L4 input strength at the three somatic gain levels.(C) Latency to the first L2/3 spike from sensory input onset as a function of L4 input strength for simulation shown in the panel B.

### Pathological E/I imbalance in the SETA circuit

Pathological excitatory/inhibitory (E/I) balance, particularly involving PV interneurons, has been proposed as a shared circuit mechanism underlying autism spectrum disorder and schizophrenia (Rubenstein and Merzenich, 2003; Lewis et al., 2005; Dienel and Lewis, 2019; Yizhar et al., 2011). However, the general E/I hypothesis does not specify where in a cortical circuit the imbalance occurs, nor does it predict why different imbalances should produce qualitatively distinct syndromes. The SETA model provides this specificity: because different interneuron types implement distinct nodes of the signed prediction error computation, E/I imbalances at different nodes produce different computational failures with different behavioural consequences. We examined two perturbations within the SETA circuit.

#### L4 excitatory drive / basket cell inhibitory balance - surprise threshold and overlearning

Under normal conditions, the modelled L4 sensory input recruits basket cells that suppress the L2/3 output, establishing the threshold for burst versus singlet firing. Reducing basket cell conductance relative to L4 excitatory drive lowers the burst firing threshold (Fig. 4A). L2/3 is consequently driven too strongly, overactivating the rest of the circuit. This translates to a higher P(BAC) (Fig. 1C) and enhanced LTP at the L5 ET apical tuft.

**Figure 4:**
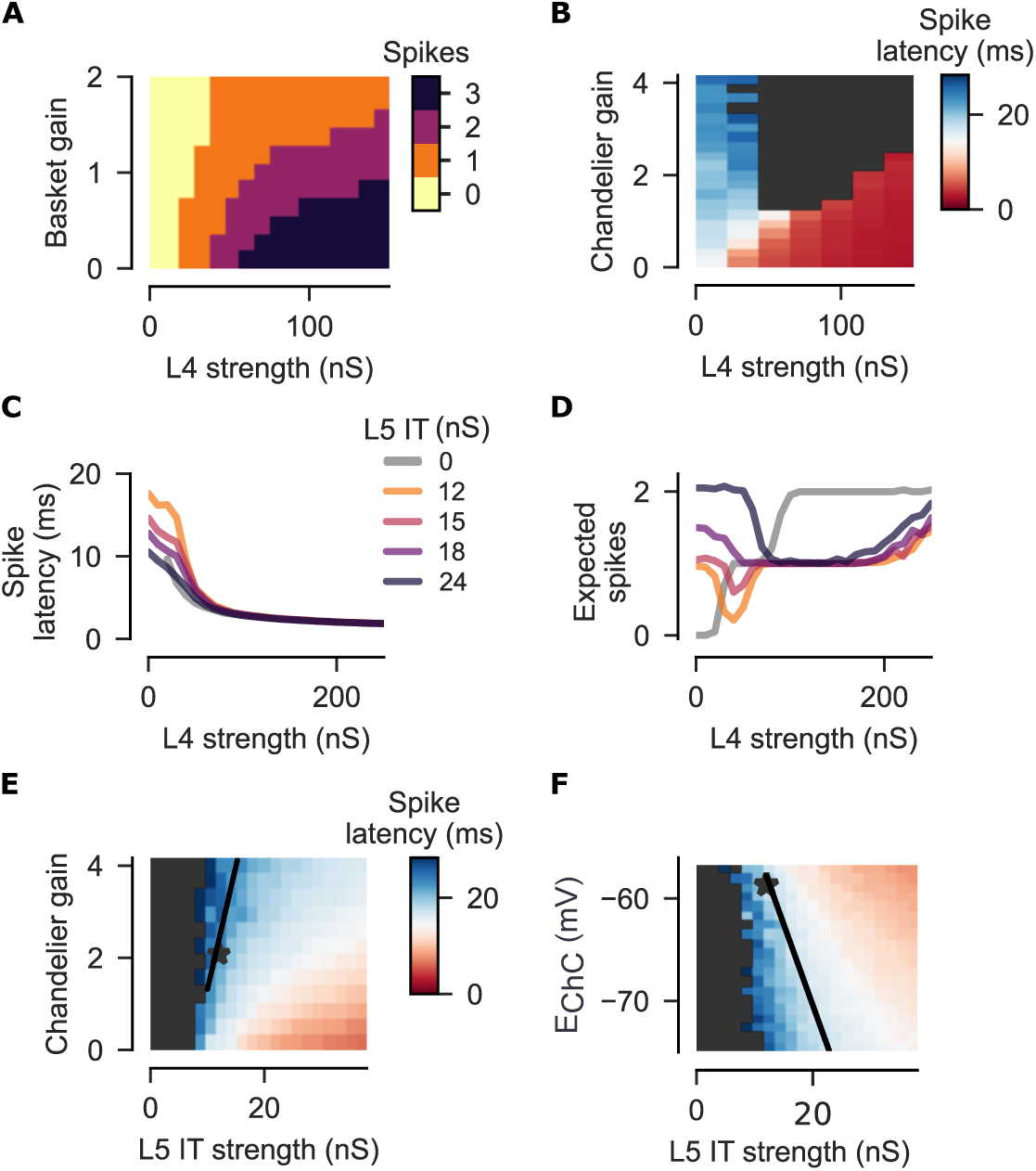
Temporal clamp failure reproduces positive psychotic symptoms. (A) Decreased basket cell conductance increases the probability of an L2/3 burst for a given sensory conductance, thereby increasing P(BAC). The plot shows L2/3 spike count as a function of Basket cell conductance and its relative scaling with L4 Excitatory cell conductance.(B) Decreased Chandelier cell conductance reduces latency to the first L2/3 spike from sensory input onset. The plot shows spike latency as a function of Chandelier cell conductance and its relative scaling with L4 Excitatory cell conductance.(C) Increased L5 IT conductance leads to shorter latency to the first L2/3 spike from sensory input onset and lowers burst threshold (D). Chandelier cell conductance was fixed at the default level (25 nS).(E) Latency to the first L2/3 spike from prediction signal onset can be maintained to match L5 IT conductance increase by scaling the chandelier gain or E_ChC_ (F). Latencies shown when sensory conductance is absent. The black lines show change required to maintain spike latency for increasing L5 IT conductance.

We propose that the reduced basket cell gain causes circuit-level sensory hypersensitivity through a compounding dual mechanism: first, the excessive burst firing amplifies sensory responses; second, the excessive bursts increase positive error signals driving additional learning. This dual mechanism may provide a biological substrate for sensory overstimulation and cognitive model overfitting (Van de Cruys et al., 2014), phenotypes of autism spectrum disorder (ASD), which are postulated to be a consequence of excitation/inhibition (E/I) imbalance and altered synaptic plasticity (Rubenstein and Merzenich, 2003; Nelson and Valakh, 2015). The reported reduction of PV basket cell density in ASD (Gogolla et al. 2009; Hashemi et al. 2017) is consistent with this specific perturbation.

#### L5 IT excitatory drive / chandelier cell inhibitory balance - sign errors and hallucinations

The second E/I perturbation concerns the balance between excitatory drive arriving at L2/3 from L5 IT prediction signals and the chandelier cell conductance implementing the temporal clamp. Reducing chandelier cell conductance relative to L5 IT excitatory drive decreased L2/3 spike latency for a given L4 strength reducing the L4 threshold required for generating a positive error (Fig. 4B). To further measure this, we next simulated how prediction error encoding changed as L5 IT conductance increased without a corresponding increase in chandelier conductance, to create an E/I imbalance. In the absence of sensory input, release-spike latency shortened with rising predictive drive (Fig. 4C), and for strong L5 IT inputs could produce a doublet rather than a singlet (Fig. 4D). Within SETA, these doublets with shortened latencies are more likely to generate sign errors: a release spike(s) arriving within the LTP rather than LTD window erroneously encodes a positive prediction error, potentiating the circuit’s association between a context and a predicted sensory event that did not occur. Overconfident prior beliefs of this kind have been proposed as the computational principle underlying hallucinations (Fletcher and Frith, 2009; Corlett et al., 2019). We also observed that weak sensory inputs interacting with strong L5 IT drive had a lower burst threshold (Fig. 4D). These bursts signal a strong positive error on weaker sensory inputs, providing a mechanistic account of aberrant salience, the inappropriate assignment of significance to neutral stimuli (Kapur, 2003), as a byproduct of the same E/I imbalance.

We simulated two parameters that could compensate for the imbalance introduced by a stronger L5 IT drive: (1) increase of the chandelier conductance and (2) a decrease of the chloride reversal potential of the chandelier synapse (E_ChC_), which shifts with maturation of the potassium-chloride cotransporter KCC2 (Rinetti-Vargas et al. 2017) (Fig. 4E–F). Either change restored the latency of the gated release spike after an increased L5 IT drive created an E/I imbalance. Positive psychotic symptoms, in this framework, may arise from failure of such compensation.

The lowered burst threshold due to stronger L5 IT drive differs fundamentally from the basket-cell-related E/I imbalance. Reduced basket cell conductance lowers the surprise threshold for any sensory input, independent of predictive state. Reduced ChC conductance relative to L5 IT drive lowers the burst threshold only in the presence of a prediction: neutral stimuli become salient specifically because a prediction has been made, not because the sensory input itself is stronger. This distinction generates a directly testable prediction: in ASD, mismatch and surprise responses should be abnormally large even in the absence of established predictions, whereas in schizophrenia the abnormality should be specifically amplified in contexts where strong predictions are held.

### In vivo evidence from Neuropixels recordings in mouse V1

The SETA model makes experimentally testable predictions about the relationship between L2/3 spiking and L5 BAC probability in vivo: positive errors should produce synergistic coincidence between the L2/3 spike and the L5 ET bAP, increasing L5 burst rate in proportion to the degree of surprise; negative errors should do the opposite, with L2/3 spikes avoiding the L5 ET LTP window and actively suppressing bursting.

To test these predictions, we analysed Neuropixels recordings from mouse primary visual cortex (V1) published by Bennett et al. (2025). The dataset was collected from mice trained in a visual change detection task, during which sequences of repeated natural visual stimuli were presented in blocks with unpredictable stimulus omissions and changes of the repeated stimulus identity (Fig. 5A). A stimulus omission should lead to a negative error due to lower sensory input than expected. The stimulus identity change could lead to negative error due to the missing predicted input, but we reasoned that in the cells with strong stimulus-evoked inputs, the influence of negative error on the spike-timing is surpassed by that of the positive error as in our simulations (Fig. 2D). To test our model predictions about increased spike bursts on positive and decreased spike bursts on negative error, we analysed the fraction of spikes occurring within bursts (FSB) in L5 neurons that we found responsive to the image stimuli (see Methods).

**Figure 5:**
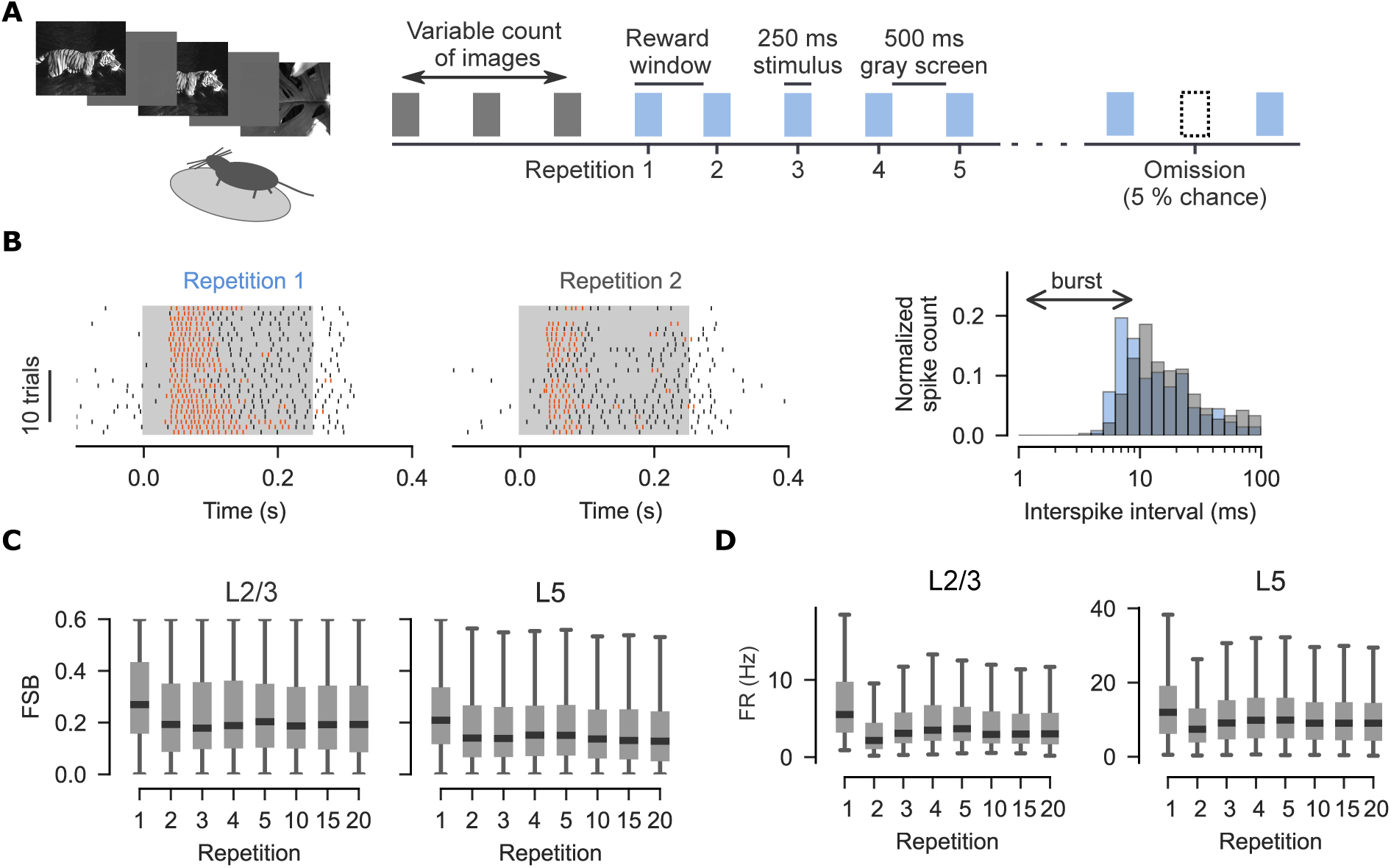
L5 bursting scales with positive errors. (A) Schematic of the image change detection task.(B) Left: Raster showing an example spiking of L5 pyramidal cell recorded in primary visual cortex in response to presentation of the same image when it follows an image change (Repetition 1) and during a subsequent presentation (Repetition 2). Spikes that are part of a burst are shown in orange. The stimulus window is marked by grey background. Right: Histogram of interspike intervals during the stimulus presentation shown in on the Left. Spike count was normalised between the Repetition 1 and 2.(C) The fraction of spikes in a burst (FSB) is highest for Repetition 1 in the L2/3 and L5 neurons and decreases with repetition in L5 neurons. The horizontal lines indicate the median, the box bounds represent the 25th and 75th percentiles, and the whiskers extend to the minimum and maximum data points. Mixed-effects model on difference from Repetition 1: for L2/3 intercept: t(1860) = -10.35, p < 10^−25^, CI = [-0.07, -0.05]; L2/3 slope: t_(1860)_ = -0.59, p = 0.55, CI = [-0.001, 0.004]; L5 intercept: t(7523) = -22.3, p < 10^−109^, CI = [-0.055, -0.046], L5 slope: t_(7523)_ = 8.1, p < 10^−15^, CI=[-0.0011,-0.0007].(D) Change in firing rate as a function of image repetition. The horizontal lines indicate the median, the box bounds represent the 25th and 75th percentiles, and the whiskers extend to the minimum and maximum data points.

#### LTP-favouring conditions: L5 burst rate scales with positive error magnitude

Consistent with the first prediction, the fraction of L5 spikes occurring within bursts (FSB) was highest for the first stimulus presentation and decreased with repeated presentations (Fig. 5B–C). Similarly, L2/3 neurons had highest burst rate for the first stimulation in line with our simulation results for unpredicted L4 input (Fig. 2A, left). For both L5 and L2/3, FSB decreased markedly from the first to second repetition. A linear mixed-effects model showed FSB decreased at a low rate with further repetitions in L5, while it remained stable in L2/3 (Fig 5C). The firing rate followed a distinct time course over the same repetitions: it fell at the second presentation and then recovered slightly to a stable level (Fig. 5D). Neither L2/3 firing rate nor L5 bursting declined to zero. This is consistent with our model, in which a perfectly predicted stimulus can produce a breakthrough spike rather than silence (Fig. 2A); a matched prediction is thus read out as a stable, non-zero baseline in both firing rate and bursting rather than as an absence of either.

The decrease in L5 bursting was reversed when a repeated presentation followed an unpredictably omitted stimulus. Post-omission stimuli drove significantly higher burst rates than pre-omission stimuli (Fig. S1A–C), consistent with increased positive error for a weaker predictability of the sequence following the unexpected omission. Burst probability also scaled with stimulus novelty: the first presentation of a newly introduced image drove higher burst rates than the first presentation of a familiar image (Fig. S1D–E). Because firing rates did not differ between novel and familiar presentations (Fig. S1F), this was specific to bursting rather than a change in overall drive. Together with the within-image repetition effect (Fig. 5C), it indicates that L5 burst rate tracks the magnitude of the positive prediction error rather than stimulus identity or firing rate per se.

#### LTD-favouring conditions: L5 burst rate is suppressed below spontaneous during omission

L5 neurons showed near-zero bursting during stimulus omission, significantly below spontaneous activity outside the stimulus blocks (Fig. 6); within SETA, this absence of bursting indicates that L2/3 input did not drive BAC firing, favouring LTD. The omission firing rate was also lower than spontaneous in both L2/3 and L5, consistent with the omission occurring with suppression (or decreased excitatory drive) for both L2/3 and L5 ET neurons.

**Figure 6:**
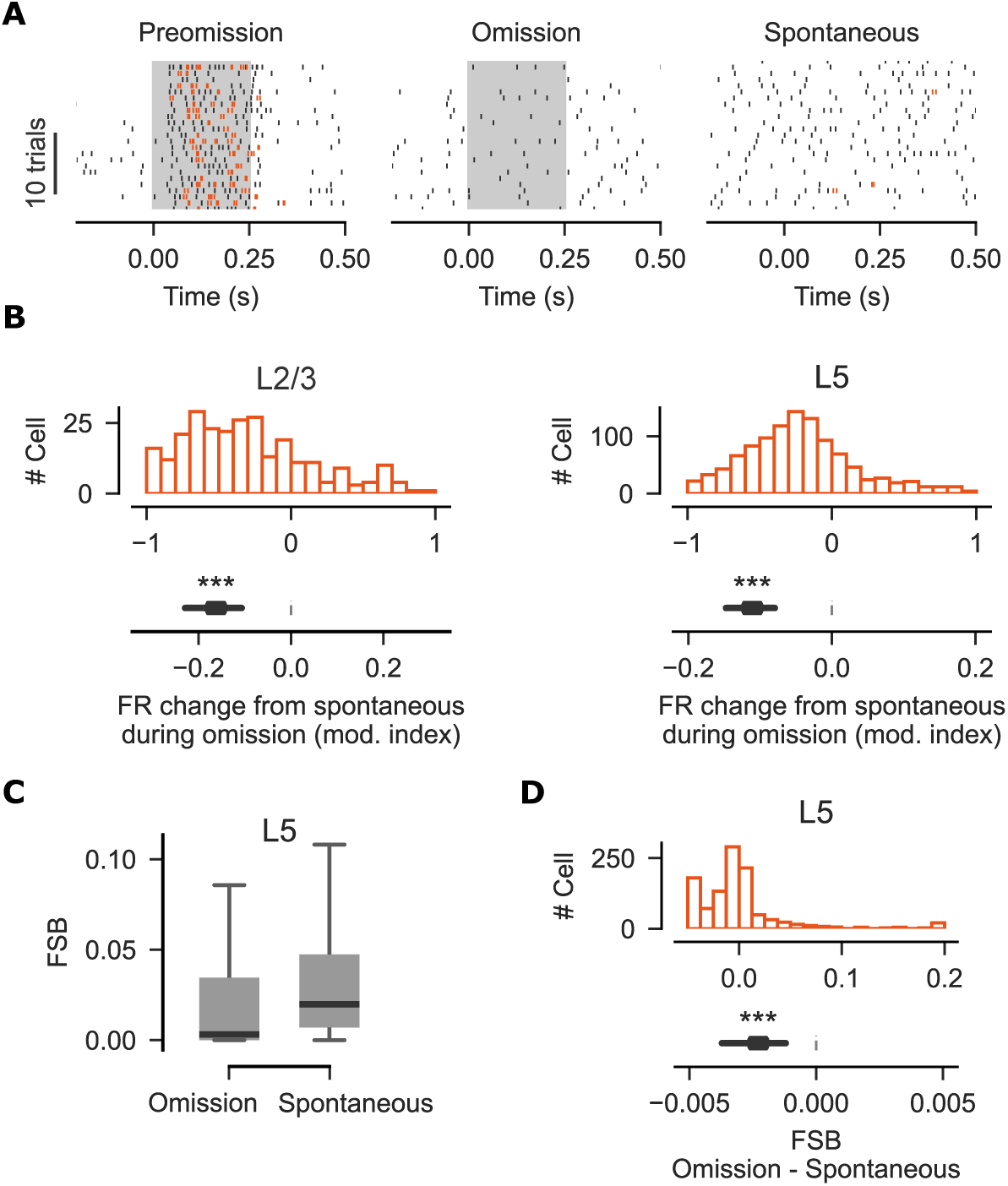
L5 bursting decreases with negative error. (A) Raster comparing spiking activity of a L5 pyramidal cell during image omission, presentation immediately before omission, and spontaneous activity. Spontaneous activity was split into 1 s trials for visualisation purposes only.(B) Top: Histogram comparing firing rate (FR) during omission to spontaneous activity periods. Bottom: Confidence interval for the change. Linear mixed-effects model L2/3: t_(265)_ = -10.4, p < 10^−24^, CI = [-0.23, -0.11], n = 266 cells from n = 78 recordings; L5: t_(1074)_ = -12.8, p < 10^−36^, CI = [-0.15, -0.08], n = 1075 cells from n = 98 recordings.(C) Barplot and (D) histogram showing decreased bursting in L5 as measured by FSB during omission compared to the spontaneous activity period. Mixed-effects model: t_(1074)_ = -7.1, p < 10^−11^, CI = [-0.004, -0.001], n = 1075 cells from n = 98 recordings.

#### L5 bursts are decoupled from L2/3 spiking on a trial-by-trial basis

We next analysed trial-by-trial noise correlations between L2/3 and L5 population activity. While overall L5 activity correlated with L2/3 activity, L5 burst rate was not predicted by either L2/3 or L5 firing rate. This held for both first repetition (Fig. S2A) and post-omission conditions (Fig. S2B). This decoupling demonstrates that L5 burst probability does not simply depend on the number of L2/3 and L5 spikes, and is consistent with our model, in which other factors, such as relative spike latency, control bursting on positive error.

#### Locomotion amplifies positive error and reduces negative-error drive

We next used locomotion as a proxy for arousal (Reimer et al. 2014; Reimer et al. 2016; Fu et al. 2014). Similar to the differential modulation of signed errors previously observed under varying stimulus uncertainty (Leonardon et al. 2025), we did not assume arousal to exert a uniform scaling of both errors. Running increased L2/3 and L5 burst rate on the first repetition of the image (Fig. 7A), consistent with our simulation of positive error gain in L2/3 (Fig. 3B) increasing with the elevated precision associated with arousal. Running also increased L2/3 and L5 firing rate (Fig. 7B). During omission, the fraction of L5 spikes occurring within bursts (FSB) did not differ between running and stationary conditions (Fig. 7C): the burst-associated fraction of spiking, and with it the sign of the error, was preserved, in spite of a reduction in firing rates during running in both L2/3 and L5 (Fig. 7D). The negative error was thus similar for running and stationary cases while positive error increased in magnitude during running.

**Figure 7:**
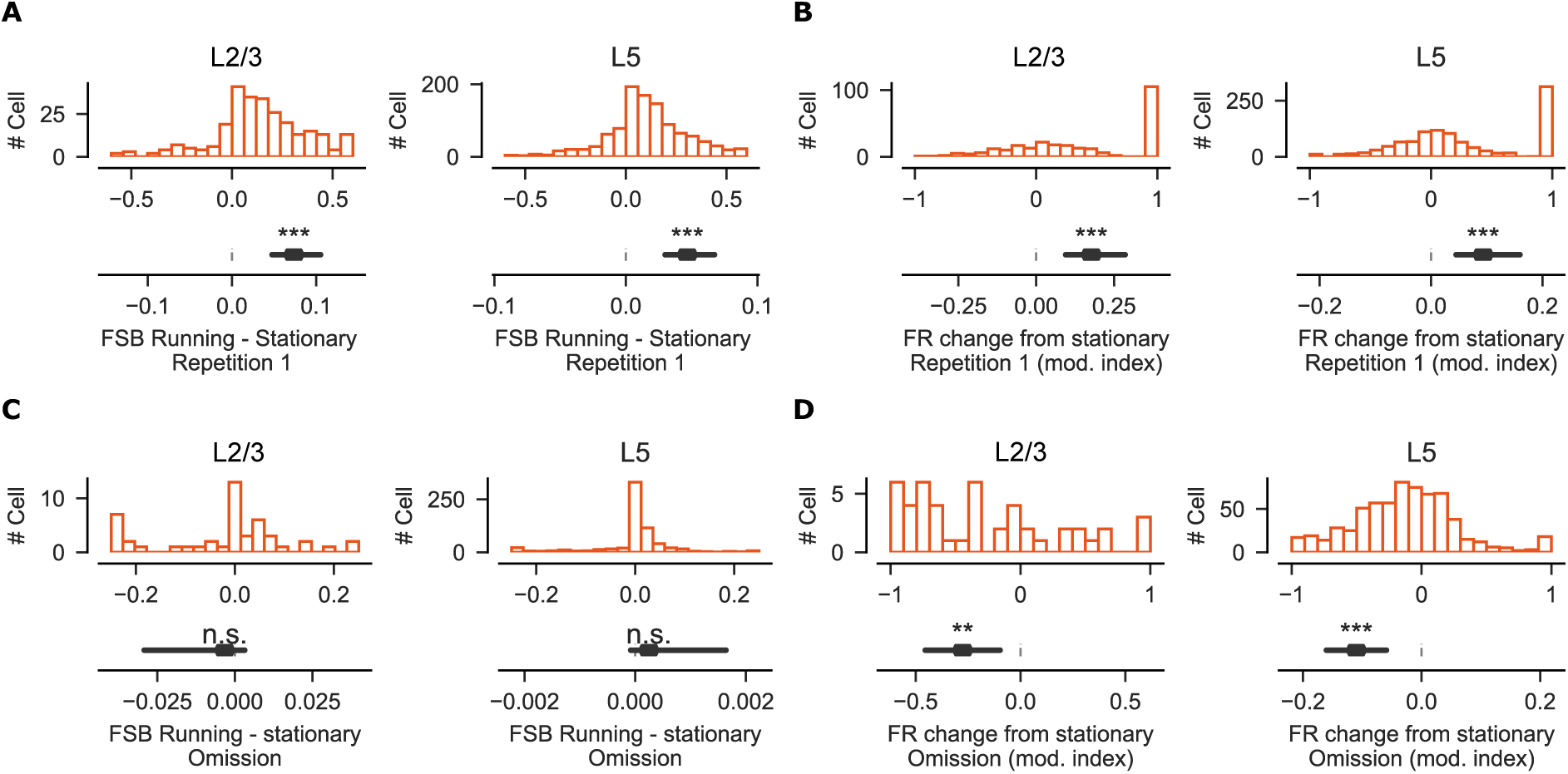
Locomotion scales positive but not negative error. (B) Top: Histogram comparing FSB during the first image repetition when mouse was running to those when mouse was stationary. Bottom: Confidence interval for the change. Mixed-effects model for L2/3: t_(265)_ = 10.1, p < 10^−23^, CI = [0.05, 0.10], n = 266 cells from n = 78 recordings; L5: t_(1074)_ = 9.7, p < 10^−21^, CI = [0.03, 0.06], n = 1075 cells from n = 98 recordings.(C) As in (A) but for change in FR. Linear mixed-effects model L2/3: t_(265)_ = 7.3, p < 10^−12^, CI = [0.09, 0.29], n = 266 cells from n = 78 recordings; L5: t_(1074)_ = 6.4, p < 10^−9^, CI = [0.05, 0.16], n = 1075 cells from n = 98 recordings.(D) As in (A) but for change in FSB during stimulus omission. Linear mixed-effects model for L2/3: t_(46)_ = -1.0, p = 0.31, CI = [-0.03, 0.003], n = 47 cells from n = 35 recordings; L5: t_(657)_ = 1.25, p = 0.21, CI = [-0.0001, 0.0016], n = 658 cells from n = 94 recordings.(E) As in (A) but for change in FR during stimulus omission. Linear mixed-effects model for L2/3: t_(46)_ = -3.1, p = 0.002, CI = [-0.45, -0.10], n = 47 cells from n = 35 recordings; L5: t_(657)_ = -4.3, p < 10^−4^, CI = [-0.16, -0.06], n = 658 cells from n = 94 recordings.

## DISCUSSION

The Signed Error by Timing Asymmetry (SETA) model provides a parsimonious solution to align the representation of signed prediction error with the requirements for plasticity underlying learning. By encoding error sign in the relative timing of L2/3 output rather than in the firing rate of dedicated populations, SETA eliminates the need for a dual-population architecture and grounds plasticity directly in the burst-dependent biophysics of L5 pyramidal neurons. The chandelier-cell temporal clamp converts a question of *whether* L2/3 fires into one of *when*, and the resulting latency, read out by the L5 ET BAC threshold, is translated into the direction of plasticity at apical synapses. Oblique synapses play a further circuit-level metaplastic role: those whose timing reliably contributes to BAC firing are strengthened, increasing the learning rate at associated tuft synapses; those that repeatedly fail to contribute to BAC firing undergo progressive LTD and eventual pruning, protecting the circuit from noisy associations. This progressive strengthening of reliable oblique synapses is consistent with burst-dependent learning frameworks in which coincident activity coordinates plasticity across hierarchical circuits (Payeur et al. 2021; Sacramento et al. 2018).

### Relationship to existing predictive coding frameworks

The canonical predictive coding model (Bastos et al. 2012) proposes that error units and representation units occupy distinct cortical layers, with bottom-up connections carrying prediction errors and top-down connections carrying predictions. SETA preserves this laminar division, sensory drive from L4, prediction error calculation in L2/3, and prediction signals emerging from deep cortical layers, but departs from canonical accounts in three respects. First, local recurrent L5 IT circuits provide the predictive signal, in contrast to inputs arriving at L1, which instead provide a contextual signal to L5 ET neurons by integrating top-down feedback from higher-order areas. This resolves the ambiguity noted by Keller and Mrsic-Flogel (2018) regarding the source of the prediction signal. Second, the negative prediction error is implemented not by Sst-mediated subtraction at the L2/3 apical dendrite (Hertäg and Clopath 2022) but by ChC-mediated temporal gating, a mechanism with the appropriate millisecond-scale dynamics. The difference is computationally important: a temporal gate acts as a precisely executed veto, while retaining some flexibility (±5 ms relative to sensory input) over the time window this veto is operational, blocking or delaying weak input and raising the threshold for doublets. Third, precision weighting (Feldman and Friston 2010) is implemented by the somatic gain of L4 and L5 IT projections, which provides a yoked excitatory drive and feedforward inhibition to L2/3 neurons- maintaining E/I balance while preserving error sign. A further implication concerns the symmetry of precision. Our simulation, following previous models, co-varies the gain on the sensory and predictive streams and co-scales positive and negative error magnitude. The in vivo data, however, show an asymmetry in scaling - locomotion amplified the positive error while L5 bursting was similarly suppressed by negative error. This is more consistent with arousal acting as a directional modulator that biases processing toward bottom-up input (Yu and Dayan 2005; Moran et al. 2013) than with a single symmetric gain and suggests that a fuller account of precision in SETA would scale the sensory and predictive streams independently rather than together.

A further departure concerns the meaning of perfect prediction. Standard predictive coding accounts treat perfect prediction as neuronal silence, and the Free Energy Principle frames learning as surprise minimisation, progressively reducing prediction error as the generative model improves (Friston 2010). SETA is consistent with this: as the prediction signal improves, L2/3 doublets, which drive BAC with higher probability and signal strong surprise, become less frequent, reserved for events that exceed prediction. Within this framework, however, silence is not a plasticity-neutral state, as it leaves apical tuft synapses biased toward LTD whenever the bAP fires, eroding learned associations. The computationally meaningful equilibrium for the synaptic weight is instead reached when a delayed L2/3 singlet at intermediate latency balances LTP and LTD over repeated trials, minimising the metabolic cost associated with bursting (Laughlin and Sejnowski 2003). We interpret the decrease of L5 ET burst rate with image repetition as bursts approaching a plasticity equilibrium, where the probabilities of potentiation and depression are matched, so that a correctly predicted, stable input produces no net synaptic change. The parallel non-zero floor in L2/3 firing rate is consistent with a sparse but non-silent response to a perfectly predicted stimulus. As the model improves, L5 ET bursting is minimised, reserving the metabolic cost of BAC firing for unexpected sensory inputs. Surprise minimisation in SETA is therefore burst minimisation rather than spike minimisation: the goal is not silence but a regime of sparse singlet firing in which BAC-driven bursts are reserved for large positive prediction errors. In this framework, L2/3 doublets update the model; singlets refine it. At the endpoint of learning, repeated LTP at tuft synapses can strengthen apical associations sufficiently that BAC firing is supported by near-coincident basal and tuft depolarisation alone; the synaptic potentiation becomes self-sustaining, no longer dependent on the oblique gating mechanism.

### Reconciling axoaxonic gating with the dual-population model

The canonical predictive coding architecture posits a functional dichotomy in L2/3, with discrete populations computing positive or negative errors (Attinger et al. 2017; Jordan & Keller 2020). Recent transcriptomic profiling supports this binary view, mapping positive and negative errors onto Rrad- and Adamts2-enriched pyramidal subtypes, respectively (O’Toole et al. 2023). However, a third Agmat-enriched population (Tasic et al. 2018; O’Toole et al. 2023) suggests a continuous transcriptomic and functional gradient rather than a strict dichotomy. SETA reframes the dual populations as two extremes of a depth-dependent gradient. Feedforward L4 sensory drive concentrates at the deep border of L2/3 and decays superficially (Weiler et al. 2023), while L5 IT and ChC inputs are distributed in the opposite direction, increasing superficially (Lefort et al. 2009; Schneider-Mizell et al. 2021). Under SETA, the error sign is not hardwired by cell type but determined by how readily a neuron overrides this depth-dependent clamp. This predicts that deep L2/3 neurons (Rrad-enriched) override the clamp rapidly to produce short-latency positive errors; superficial neurons (Adamts2-enriched) are reliably clamped, producing delayed negative errors. The precise makeup of these three populations for L2/3 projections, both to L5 ET basal and oblique dendrites, and their implications for gating plasticity, remains to be tested.

### E/I imbalance and psychiatric disorders

Pathological E/I imbalance involving PV interneurons has been proposed as a shared mechanism underlying both autism spectrum disorder and schizophrenia (Rubenstein and Merzenich 2003; Lewis et al. 2005). SETA specifies that the location of the imbalance in the prediction error circuit produces qualitatively distinct failures: basket cell hypofunction lowers the surprise threshold without corrupting error sign, producing sensory hypersensitivity; ChC hypofunction causes sign errors corrupting the direction of learning, leading to hallucinations. The node of E/I disruption, rather than the disruption itself, determines the clinical consequence.

The schizophrenia case is notable because multiple independent molecular routes converge on the same computational lesion: insufficient chandelier conductance relative to L5 IT prediction drive. NMDA receptor hypofunction in PV interneurons (Singh et al. 2022; Krystal et al. 1994; Homayoun and Moghaddam 2007; Lahti et al. 1995) reduces the probability that chandelier cells reach firing threshold in response to L5 IT prediction. Reduced GAD67 expression in PV boutons (Hashimoto et al. 2003; Lewis et al. 2005) reduces conductance per clamp event. AIS scaffolding loss (ANK3; Tai et al. 2019) destabilises postsynaptic GABA_A_ clustering. Chloride transporter dysregulation (Hyde et al. 2011; Arion and Lewis 2011) collapses the gradient that allows the clamp to strengthen with maturing predictions. Compensatory upregulation of postsynaptic α2 GABA_A_ receptors (Volk et al. 2002) may reflect attempts by the brain to preserve clamp function as E/I balance deteriorates.

Extending our framework further, we speculate that hallucinations could arise through diverse pharmacological or physiological routes, provided they converge on the same computational endpoint: an abnormally elevated P(BAC). This could occur in at least four ways: (1) weakening of the temporal clamp, advancing the release spike; (2) raised apical-tuft excitability, lowering the threshold for BAC firing; (3) excessive predictive drive; and (4) lengthening of the LTP window. We propose that these converge on a single failure mode: during over-prediction, a higher P(BAC) causes sign errors in L5 ET neurons. This provides a unifying framework for how diverse biotypes and pharmacological states can converge to generate hallucinations, and explains why the search for a single, universal molecular cause of psychosis has proven elusive (Singh et al., 2022; Trubetskoy et al., 2022).

### Experimental validation of the model

Our model posits that encoding of the error sign is controlled by L2/3 spike timing relative to the L5 ET bAP. When the L2/3 input arrives in a short delay after the bAP, it promotes BAC and associated somatic bursting. Consistent with this, our analysis demonstrates that the L5 burst rate scales with the magnitude of positive error. While the burst rate correlates with firing rate, trial-by-trial analysis of noise-correlations reveals that burst occurrence is decoupled from population firing rate. This independence is consistent with our explanation that prediction error depends on sub-millisecond spike-timing and dendritic integration within L5 ET neurons.

Unlike positive errors anchored to an external stimulus, a negative error during omission relies entirely on an internal prediction. The lack of anchoring to an external stimulus means that the exact timing of L5 IT prediction signal spikes varies from trial to trial. Consequently, the timing of the delayed L2/3 spike also varies with respect to the omission onset. Nevertheless, we find that L5 bursts are suppressed during negative errors compared to the spontaneous level. The change is compatible with our model, which posits that low BAC probability biases plasticity towards LTD, encoding negative error.

The SETA model proposes that the same L5 ET population multiplexes positive and negative error, we restricted our analysis to image-responsive neurons, as these will necessarily include the neurons that burst-fire in response to the stimulus, as required for representing positive error. This approach diverges from unrestricted whole-population analyses of the same dataset, which reports a gradual ramping of activity during omissions in a small fraction of visual cortex neurons (Nitzan & Buzsáki, 2025). Importantly, ramping activity can be driven by mechanisms other than predictive inputs; it can reflect top-down modulation, shifts in neuromodulatory tone, or intrinsic neuronal properties such as recovery from adaptation.

### Limitations and future directions

Several limitations merit explicit acknowledgement. First, although the biophysical conditions for the gated release spike are established in neocortical L2/3 (Woodruff et al. 2009, 2011), and paired recordings in basolateral amygdala demonstrate axo-axonic-mediated spike delays consistent with the SETA LTD window (Veres et al. 2014), the gated release spike has not been directly recorded in neocortical L2/3 in the predictive coding circuit context SETA requires. The BLA recordings were obtained under hyperpolarising axo-axonic conditions, meaning the delay mechanism is analogous but not identical to the neocortical clamp proposed here. Importantly, our model produces equivalent timing under both depolarising and hyperpolarising clamps, parametrically tuneable via L5 IT input strength and chandelier conductance. Direct confirmation would require paired recordings from L5 IT, chandelier, and L2/3 pyramidal neurons during prediction and omission.

The second limitation is that the Allen Institute dataset does not permit definitive separation of L5 ET from L5 IT neurons; while L5 IT neurons show close to zero bursting, definitive cell-type-specific testing requires antidromic stimulation or transgenic labelling. Future Neuropixels recordings combined with optogenetic identification of L5 ET neurons (Economo et al. 2018) would allow direct testing of the LTP/LTD prediction at the specific cell type. A related limitation is that locomotion is a global state variable that co-modulates many interneuron types in heterogeneous directions (Bugeon et al. 2022); it therefore cannot dissociate the precision of the sensory and predictive streams, and we use it only as a proxy for elevated precision.

Finally, SETA does not yet specify the learning dynamics of the L5 IT generative model. Extending the present framework to include the role of L2/3 prediction errors in modifying L5 IT recurrent networks, and the propagation of L2/3 prediction errors to higher order cortical areas is the necessary next step toward a complete computational account of how the generative model is updated over experience.

## Author Contributions

**D.B.** Conceptualization, Methodology, Funding Acquisition, Writing – Original Draft, Writing – Review and Editing. **P.J.:** Methodology, Software, Formal Analysis, Data Curation, Visualization, Funding Acquisition, Writing – Review and Editing.

## Acknowledgements

We thank Shohei Furutachi and Maxwell Shinn for comments on the manuscript.

## Funding

RNID Fellowship: FG24/9 (P.J.), Biotechnology and Biological Sciences Research Council (BBSRC) Research Grants: BB/Z51665X/1, BB/Y010345/1 (D.B.).

## Code Availability

Simulation and data analysis codes will be made available on publication.

## Declaration of AI-assistance

During the preparation of this work, the authors used AI tools (Claude/Gemini) for conceptual discussions, assisted coding, and manuscript edits. The authors take full responsibility for the final content of the manuscript.

## STAR METHODS

### Key resource table

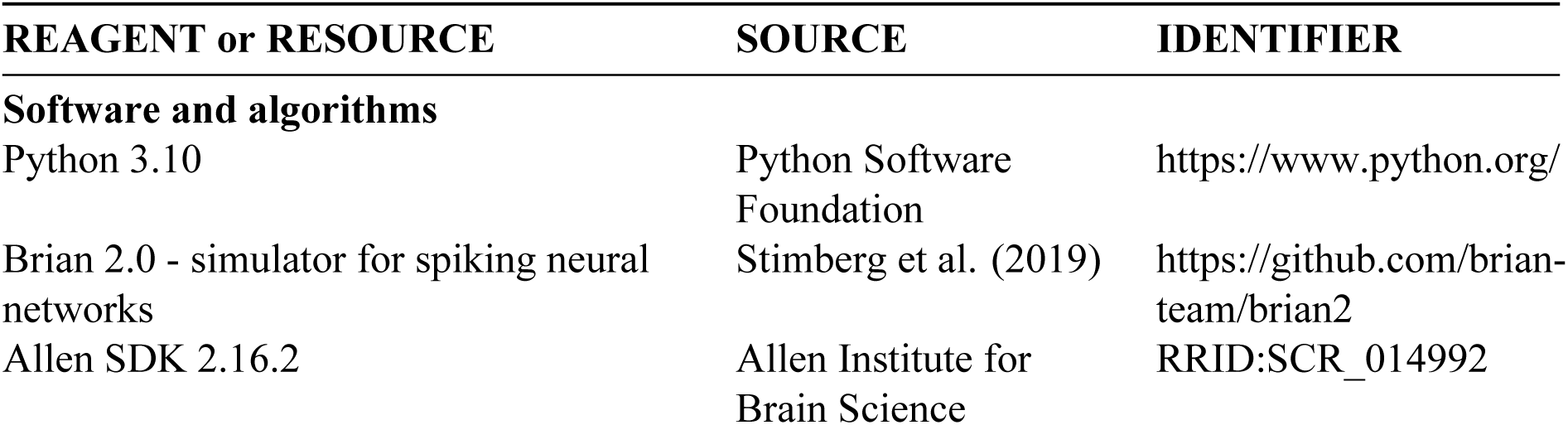
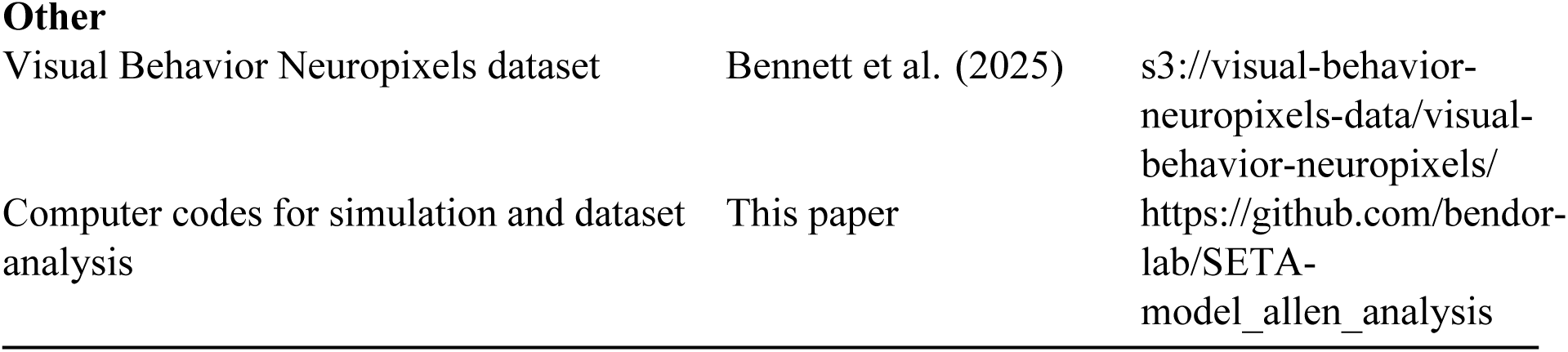

### Method details

#### Computational model of L2/3 neuron

The simulated layer 2/3 (L2/3) pyramidal neuron was modelled using a two-compartment leaky integrate-and-fire (LIF) model separating somatic integration from axonal spike initiation. The two compartments represent the soma and the axon initial segment (AIS), coupled by an axial conductance g_a_. The temporal evolution of somatic voltage (V_soma_) and AIS voltage (V_ais_) is governed by:

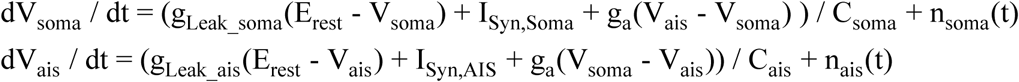

I_Syn,Soma_ and I_Syn,AIS_ represent synaptic currents of the soma and AIS, respectively. Somatic membrane capacitance C_soma_ = 0.281 nF, time constant 30 ms, and leak conductance g_Leak_soma_ = 9.4 nS; AIS capacitance C_ais_ = 0.0145 nF (5% of somatic to account for their relative sizes), time constant 11 ms and leak conductance g_Leak_ais_ = 1.25 nS. Both compartments share resting potential of E_rest_ = -70.0 mV. Axial coupling conductance g_a_ = 18.0 nS. Spike generation occurs exclusively at the AIS upon V_ais_ reaching threshold -50.0 mV, after which both compartments reset to -70.0 mV for a hard refractory period of 2.0 ms.

Voltage updates include an additive Gaussian noise term implemented as n(t) = σ / sqrt(τ_noise_) * xi(t), where xi ∼ N(0, 1), σ = 3 mV, τ_noise_ = 10 ms. Each simulation was repeated 80 times to average the effects of the Gaussian noise, with the exception of results presented in Fig. 4A,B,E,F where it was repeated 10 times for simulation efficiency.

The simulations were run using Brian 2.0 simulator package (Stimberg et al. 2019).

##### Synaptic connectivity and kinetics

Four synaptic input streams are distributed across the two compartments. All conductances are modelled with alpha functions: g(t) = g_max_ * (t / τ) * exp(1 – t / τ) and signify total input provided from a given cell class parametrised by its maximal conductance g_max_ and time constant τ.

##### Somatic inputs

Layer 4 (L4) excitatory cells sensory input. Fast AMPAergic feedforward excitation. Time constant of 4.0 ms; reversal potential of 0 mV. Conductance was set to 60 nS to generate a single spike response of the L2/3 pyramidal neuron.

Layer 5 (L5) IT input. Time constant of 8.0 ms; reversal potential 0 mV. Default conductance was set to 12 nS.

Basket Cell input. Time constant of 6.0 ms; reversal potential -75.0 mV. Basket conductance was set as a scaled L4 sensory conductance (1.2x by default) instantiating strong feedforward inhibition. Conductance onset was delayed by 2.5 ms relative to its driving L4 excitatory cell input.

##### Axonal Input

Chandelier Cell (ChC) input. Targeted inhibition to the AIS compartment only. Time constant of 4.0 ms; reversal potential of -58.0 mV. This depolarised reversal (between resting -70 mV and threshold -50 mV) acts as a voltage clamp on the AIS, suppressing spike initiation without classical hyperpolarisation. Default clamping conductance was set to 25 nS and the conductance onset was delayed by 1.5 ms relative to its driving L5 IT input.

Scaling of L2/3 inputs by somatic gain of L4 and L5 IT inputs

Different levels of somatic gain were simulated by proportional scaling of excitatory and inhibitory L2/3 inputs. The conductance of L4, Basket, L5 IT and ChC inputs was scaled by the same multiplier: 1.0 for High, 0.5 for Mid and 0.2 for Low gain level. This multiplier represents the stimulus-evoked feedforward inhibitory gain yoked to L4 and L5 IT drive, rather than the tonic firing rate of the interneurons, which need not change in the same direction with arousal (Bugeon et al. 2022).

#### Allen Brain Observatory data analysis

To assess predictions of our model, we analysed the Visual Behavior Neuropixels dataset published by the Allen Brain Institute (resource ID: s3://visual-behavior-neuropixels-data/visual-behavior-neuropixels/, https://doi.org/10.48324/dandi.000713/0.240702.1725). The dataset contains electrophysiological recordings performed in mice with Neuropixel extracellular probes and associated behavioural data. Data were collected from mice trained in a visual change detection task. The training, task, and detailed methods for data acquisition and preprocessing were described in Bennett et al. (2025). Here, we performed analysis using spike data from channels identified in the dataset as located in the primary visual cortex (VISp).

In the first part of the recording session, mice were exposed to a series of natural image presentations on the screen, organised in blocks of repeating stimuli. The 250 ms presentations of the stimuli were interleaved with a 500 ms presentation of a grey screen. The image identity was changed after a random number of repetitions. Mice were rewarded with water for correctly licking following the image change.

We analysed the two periods of the recording session: active part of the task, during which mice were rewarded for change detection; and the period of mouse inactivity immediately following the task, to assess spontaneous neuronal activity.

To assess the effect of image novelty on activity, we compared neuronal responses to the first image repetition in a sequence following image identity change when it was one of the first 10 blocks with that image and when it was a block repeating the image for the twentieth or later time. For this analysis, we only considered sessions when the image set was presented for the first time.

##### Locomotion detection

We classified the running behaviour during individual stimulus presentations as ‘stationary’ or ‘locomotion’ based on locomotion speed. The presentation was classified as ‘locomotion’ if the mean speed exceeded 5 cm/s, and it exceeded 3 cm/s during 75% of the presentation duration. It was classified as ‘stationary’ if the mean speed was below 5 cm/s and the speed was below 3 cm/s during 75% of the presentation duration.

##### Laminar classification

The dataset registers unit locations in the brain to the Allen Institute Common Coordinate Framework (CCFv3). We leveraged the unit’s coordinates to extract layer labels for each cortical unit (L1, L2/3, L4, L5, or L6) based on the average brain template in CCFv3, as described by Siegle et al. (2021). <u>Unit inclusion</u>

Spiking units were included in the analysis only if they satisfied the following criteria: (1) inter-spike-interval violations (Hill et al. 2011) not exceeding 0.5 ; (2) amplitude distribution cut-off ≤ 10%; (3) presence ratio ≥ 90%; (4) signal to noise ratio > 1; (5) firing rate > 0.2 Hz; (6) they were classed as ‘good’ in the published dataset.

We further restricted the units to those that were responsive to the image stimuli, based on their PSTH calculated across presentations of different images. We compared the activity during the 250 ms following the stimulus onset with that of the same duration immediately before. A unit was classified as responsive if its firing rate was significantly higher during the presentation (p-value < 0.05, Wilcoxon paired test) and it had spikes in response to ≥ 20% stimulus presentations.

When analysing joint population dynamics in L2/3 and L5 and their noise correlations, we only included trials from sessions with at least 10 putative pyramidal cells per layer. Additionally, noise correlations on the L5 FSB only included trials with spikes.

##### Identification of pyramidal neurons

The spiking analysis was performed on putative pyramidal L2/3 and L5 neurons. Candidate pyramidal neurons were first identified based on the waveform duration (peak-to-trough longer than 0.4 ms) and firing rate (lower than 10 Hz). The candidate cells were further filtered by removing putative inhibitory neurons identified in the optotagging data. The optotagging experiment used VIP-IRES-Cre;Ai32 or Sst-IRES-Cre;Ai32 transgenic mouse lines to identify Vip and Sst cells, respectively. A cell was considered to be of that specific type if optogenetic stimulation resulted in its activity exceeding 50 Hz and a z-score exceeding 2 compared to its baseline firing during the 1-10 ms window following stimulation onset.

##### Spiking metrics and burst detection

Neuronal spiking times were aligned to stimulus presentation and omission onsets, and we calculated spiking metrics over 250-ms-long periods. A burst was detected whenever two spikes had an inter-spike interval ≤ 10 ms. Next, we calculated the fraction of all spikes during the stimulus periods that were part of the detected bursts (FSB).

To analyse the change in the neuron’s firing rate (FR) from a control condition, we calculated modulation index as: (FR’ – FR_control_) / (FR’ + FR_control_).

Noise correlations were calculated between L2/3 and L5 activity and bursting to assess their trial-to-trial co-variability. First the cell’s mean response to a stimulation image at a given repetition ordinal was subtracted from the trial value. Next, population mean was calculated for the trial on neurons recorded in the same session from the same layer of primary visual cortex.

### Quantification and statistical analysis

Mixed-effects models were used for statistical analysis. The effects were assessed with a linear model or a model with the response variable transformed by the square root to satisfy the assumption of normality of residuals. The model specification included a random-effect for session to account for correlations in the activity of cells recorded in the same session. Statistical model on data in Fig. 5C modelled the fixed effects of repetition. The other figures that tested for changes in cells’ bursting or firing report the significance of the intercept. Confidence intervals for the effects were estimated from t-distribution based on the calculated t-statistic and degrees of freedom of the model.

p-values < 0.05 were considered significant. Confidence intervals are reported at the 95% level. Statistical details can be found in figure legends.

## Notes

### Competing Interest Statement

The authors have declared no competing interest.

## REFERENCES

1. Arion, D., and Lewis, D. A. (2011). Altered expression of regulators of the cortical chloride transporters NKCC1 and KCC2 in schizophrenia. Archives of general psychiatry, 68(1), 21–31.

2. Attinger, A., Wang, B., and Keller, G. B. (2017). Visuomotor coupling shapes the functional development of mouse visual cortex. Cell, 169(7), 1291–1302.

3. Bastos, A. M., Lundqvist, M., Waite, A. S., Kopell, N., and Miller, E. K. (2020). Layer and rhythm specificity for predictive routing. Proceedings of the National Academy of Sciences, 117(49), 31459–31469.

4. Bastos, A. M., Usrey, W. M., Adams, R. A., Mangun, G. R., Fries, P., and Friston, K. J. (2012). Canonical microcircuits for predictive coding. Neuron, 76(4), 695–711.

5. Bennett, C., Gale, S., Heller, G., Ramirez, T., Belski, H., Piet, A., … and Olsen, S. R. (2025). Map of spiking activity underlying change detection in the mouse visual system. bioRxiv, 2025-10.

6. Brown, S. P., and Hestrin, S. (2009). Intracortical circuits of pyramidal neurons reflect their long-range axonal targets. Nature, 457(7233), 1133–1136.

7. Bugeon, S., Duffield, J., Dipoppa, M., Ritoux, A., Prankerd, I., Nicoloutsopoulos, D., … & Harris, K. D. (2022). A transcriptomic axis predicts state modulation of cortical interneurons. Nature, 607(7918), 330–338.

8. Constantinople, C. M., and Bruno, R. M. (2013). Deep cortical layers are activated directly by thalamus. Science, 340(6140), 1591–1594.

9. Corlett, P. R., Horga, G., Fletcher, P. C., Alderson-Day, B., Schmack, K., and Powers, A. R. (2019). Hallucinations and strong priors. Trends in cognitive sciences, 23(2), 114–127.

10. Dienel, S. J., and Lewis, D. A. (2019). Alterations in cortical interneurons and cognitive function in schizophrenia. Neurobiology of disease, 131, 104208.

11. Economo, M. N., Viswanathan, S., Tasic, B., Bas, E., Winnubst, J., Menon, V., … and Svoboda, K. (2018). Distinct descending motor cortex pathways and their roles in movement. Nature, 563(7729), 79–84.

12. Feldman, H., and Friston, K. J. (2010). Attention, uncertainty, and free-energy. Frontiers in human neuroscience, 4, 215.

13. Fletcher, P. C., and Frith, C. D. (2009). Perceiving is believing: a Bayesian approach to explaining the positive symptoms of schizophrenia. Nature Reviews Neuroscience, 10(1), 48–58.

14. Friston, K. (2005). A theory of cortical responses. Philosophical transactions of the Royal Society B: Biological sciences, 360(1456), 815–836.

15. Friston, K. (2010). The free-energy principle: a unified brain theory?. Nature reviews neuroscience, 11(2), 127–138.

16. Fu, Y., Tucciarone, J. M., Espinosa, J. S., Sheng, N., Darcy, D. P., Nicoll, R. A., … and Stryker, M. P. (2014). A cortical circuit for gain control by behavioral state. Cell, 156(6), 1139–1152.

17. Gogolla, N., LeBlanc, J. J., Quast, K. B., Südhof, T. C., Fagiolini, M., and Hensch, T. K. (2009). Common circuit defect of excitatory-inhibitory balance in mouse models of autism. Journal of neurodevelopmental disorders, 1(2), 172–181.

18. Graupner, M., and Brunel, N. (2012). Calcium-based plasticity model explains sensitivity of synaptic changes to spike pattern, rate, and dendritic location. Proceedings of the national academy of sciences, 109(10), 3991–3996.

19. Hage, T. A., Bosma-Moody, A., Baker, C. A., Kratz, M. B., Campagnola, L., Jarsky, T., … and Murphy, G. J. (2022). Synaptic connectivity to L2/3 of primary visual cortex measured by two-photon optogenetic stimulation. Elife, 11, e71103.

20. Harris, K. D., and Shepherd, G. M. (2015). The neocortical circuit: themes and variations. Nature neuroscience, 18(2), 170–181.

21. Hashemi, E., Ariza, J., Rogers, H., Noctor, S. C., and Martínez-Cerdeño, V. (2017). The number of parvalbumin-expressing interneurons is decreased in the prefrontal cortex in autism. Cerebral cortex, 27(3), 1931–1943.

22. Hashimoto, T., Volk, D. W., Eggan, S. M., Mirnics, K., Pierri, J. N., Sun, Z., … and Lewis, D. A. (2003). Gene expression deficits in a subclass of GABA neurons in the prefrontal cortex of subjects with schizophrenia. Journal of Neuroscience, 23(15), 6315–6326.

23. Heindorf, M., & Keller, G. B. (2024). Antipsychotic drugs selectively decorrelate long-range interactions in deep cortical layers. Elife, 12, RP86805.

24. Helmstaedter, M., Staiger, J. F., Sakmann, B., & Feldmeyer, D. (2008). Efficient recruitment of layer 2/3 interneurons by layer 4 input in single columns of rat somatosensory cortex. Journal of Neuroscience, 28(33), 8273–8284.

25. Hertäg, L., and Clopath, C. (2022). Prediction-error neurons in circuits with multiple neuron types: Formation, refinement, and functional implications. Proceedings of the National Academy of Sciences, 119(13), e2115699119.

26. Hill, D. N., Mehta, S. B., and Kleinfeld, D. (2011). Quality metrics to accompany spike sorting of extracellular signals. Journal of Neuroscience, 31(24), 8699–8705.

27. Homayoun, H., and Moghaddam, B. (2007). NMDA receptor hypofunction produces opposite effects on prefrontal cortex interneurons and pyramidal neurons. Journal of Neuroscience, 27(43), 11496–11500.

28. Hyde, T. M., Lipska, B. K., Ali, T., Mathew, S. V., Law, A. J., Metitiri, O. E., … and Kleinman, J. E. (2011). Expression of GABA signaling molecules KCC2, NKCC1, and GAD1 in cortical development and schizophrenia. Journal of Neuroscience, 31(30), 11088–11095.

29. Jordan, R., and Keller, G. B. (2020). Opposing influence of top-down and bottom-up input on excitatory layer 2/3 neurons in mouse primary visual cortex. Neuron, 108(6), 1194–1206.

30. Kapur, S. (2003). Psychosis as a state of aberrant salience: a framework linking biology, phenomenology, and pharmacology in schizophrenia. American journal of Psychiatry, 160(1), 13–23.

31. Keller, G. B., Bonhoeffer, T., and Hübener, M. (2012). Sensorimotor mismatch signals in primary visual cortex of the behaving mouse. Neuron, 74(5), 809–815.

32. Keller, G. B., and Mrsic-Flogel, T. D. (2018). Predictive processing: a canonical cortical computation. Neuron, 100(2), 424–435.

33. Kim, E. J., Juavinett, A. L., Kyubwa, E. M., Jacobs, M. W., and Callaway, E. M. (2015). Three types of cortical layer 5 neurons that differ in brain-wide connectivity and function. Neuron, 88(6), 1253–1267.

34. Krystal, J. H., Karper, L. P., Seibyl, J. P., Freeman, G. K., Delaney, R., Bremner, J. D., … and Charney, D. S. (1994). Subanesthetic effects of the noncompetitive NMDA antagonist, ketamine, in humans: psychotomimetic, perceptual, cognitive, and neuroendocrine responses. Archives of general psychiatry, 51(3), 199–214.

35. Lahti, A. C., Koffel, B., LaPorte, D., and Tamminga, C. A. (1995). Subanesthetic doses of ketamine stimulate psychosis in schizophrenia. Neuropsychopharmacology, 13(1), 9–19.

36. Larkum, M. (2013). A cellular mechanism for cortical associations: an organizing principle for the cerebral cortex. Trends in neurosciences, 36(3), 141–151.

37. Larkum, M. E., Zhu, J. J., and Sakmann, B. (1999). A new cellular mechanism for coupling inputs arriving at different cortical layers. Nature, 398(6725), 338–341.

38. Larkum, M. E., Zhu, J. J., and Sakmann, B. (2001). Dendritic mechanisms underlying the coupling of the dendritic with the axonal action potential initiation zone of adult rat layer 5 pyramidal neurons. The Journal of physiology, 533(2), 447–466.

39. Laughlin, S. B., and Sejnowski, T. J. (2003). Communication in neuronal networks. Science, 301(5641), 1870–1874.

40. Lefort, S., Tomm, C., Sarria, J. C. F., and Petersen, C. C. (2009). The excitatory neuronal network of the C2 barrel column in mouse primary somatosensory cortex. Neuron, 61(2), 301–316.

41. Leonardon, B., Kasavica, S., Raltschev, C., Senn, W., Wilmes, K. A., & Sachidhanandam, S. (2025). Differential modulation of positive and negative prediction errors by stimulus variability in the mouse posterior parietal cortex. Communications Biology, 8(1), 1397.

42. Letzkus, J. J., Kampa, B. M., and Stuart, G. J. (2006). Learning rules for spike timing-dependent plasticity depend on dendritic synapse location. Journal of Neuroscience, 26(41), 10420–10429.

43. Lewis, D. A., Hashimoto, T., and Volk, D. W. (2005). Cortical inhibitory neurons and schizophrenia. Nature Reviews Neuroscience, 6(4), 312–324.

44. Moran, R. J., Campo, P., Symmonds, M., Stephan, K. E., Dolan, R. J., & Friston, K. J. (2013). Free energy, precision and learning: the role of cholinergic neuromodulation. Journal of Neuroscience, 33(19), 8227–8236.

45. Nelson, S. B., & Valakh, V. (2015). Excitatory/inhibitory balance and circuit homeostasis in autism spectrum disorders. Neuron, 87(4), 684–698.

46. Nitzan, N., & Buzsáki, G. (2025). Diversity of omission responses to visual images across brain-wide regions. Science Advances, 11(21), eadv5651.

47. O’Toole, S. M., Oyibo, H. K., and Keller, G. B. (2023). Molecularly targetable cell types in mouse visual cortex have distinguishable prediction error responses. Neuron, 111(18), 2918–2928.

48. Pala, A., and Petersen, C. C. (2015). In vivo measurement of cell-type-specific synaptic connectivity and synaptic transmission in layer 2/3 mouse barrel cortex. Neuron, 85(1), 68–75.

49. Paulsen, O., and Sejnowski, T. J. (2000). Natural patterns of activity and long-term synaptic plasticity. Current opinion in neurobiology, 10(2), 172–180.

50. Payeur, A., Guerguiev, J., Zenke, F., Richards, B. A., and Naud, R. (2021). Burst-dependent synaptic plasticity can coordinate learning in hierarchical circuits. Nature neuroscience, 24(7), 1010–1019.

51. Petreanu, L., Mao, T., Sternson, S. M., and Svoboda, K. (2009). The subcellular organization of neocortical excitatory connections. Nature, 457(7233), 1142–1145.

52. Pfeffer, C. K., Xue, M., He, M., Huang, Z. J., and Scanziani, M. (2013). Inhibition of inhibition in visual cortex: the logic of connections between molecularly distinct interneurons. Nature neuroscience, 16(8), 1068–1076.

53. Pi, H. J., Hangya, B., Kvitsiani, D., Sanders, J. I., Huang, Z. J., and Kepecs, A. (2013). Cortical interneurons that specialize in disinhibitory control. Nature, 503(7477), 521–524.

54. Polack, P. O., Friedman, J., and Golshani, P. (2013). Cellular mechanisms of brain state–dependent gain modulation in visual cortex. Nature neuroscience, 16(9), 1331–1339.

55. Rao, R. P., and Ballard, D. H. (1999). Predictive coding in the visual cortex: a functional interpretation of some extra-classical receptive-field effects. Nature neuroscience, 2(1), 79–87.

56. Reimer, J., Froudarakis, E., Cadwell, C. R., Yatsenko, D., Denfield, G. H., and Tolias, A. S. (2014). Pupil fluctuations track fast switching of cortical states during quiet wakefulness. neuron, 84(2), 355–362.

57. Reimer, J., McGinley, M. J., Liu, Y., Rodenkirch, C., Wang, Q., McCormick, D. A., and Tolias, A. S. (2016). Pupil fluctuations track rapid changes in adrenergic and cholinergic activity in cortex. Nature communications, 7(1), 13289.

58. Rinetti-Vargas, G., Phamluong, K., Ron, D., and Bender, K. J. (2017). Periadolescent maturation of GABAergic hyperpolarization at the axon initial segment. Cell reports, 20(1), 21–29.

59. Rubenstein, J. L., and Merzenich, M. M. (2003). Model of autism: increased ratio of excitation/inhibition in key neural systems. Genes, Brain and Behavior, 2(5), 255–267.

60. Sacramento, J., Ponte Costa, R., Bengio, Y., and Senn, W. (2018). Dendritic cortical microcircuits approximate the backpropagation algorithm. Advances in neural information processing systems, 31.

61. Sakata, S., and Harris, K. D. (2009). Laminar structure of spontaneous and sensory-evoked population activity in auditory cortex. Neuron, 64(3), 404–418.

62. Schaefer, A. T., Larkum, M. E., Sakmann, B., and Roth, A. (2003). Coincidence detection in pyramidal neurons is tuned by their dendritic branching pattern. Journal of neurophysiology, 89(6), 3143–3154.

63. Schneider-Mizell, C. M., Bodor, A. L., Collman, F., Brittain, D., Bleckert, A., Dorkenwald, S., … and Costa, N. M. D. (2021). Structure and function of axo-axonic inhibition. Elife, 10, e73783.

64. Seignette, K., Jamann, N., Papale, P., Terra, H., Porneso, R. O., de Kraker, L., … and Levelt, C. N. (2024). Experience shapes chandelier cell function and structure in the visual cortex. Elife, 12, RP91153.

65. Siegle, J. H., Jia, X., Durand, S., Gale, S., Bennett, C., Graddis, N., … and Koch, C. (2021). Survey of spiking in the mouse visual system reveals functional hierarchy. Nature, 592(7852), 86–92.

66. Singh, T., Poterba, T., Curtis, D., Akil, H., Al Eissa, M., Barchas, J. D., … and Daly, M. J. (2022). Rare coding variants in ten genes confer substantial risk for schizophrenia. Nature, 604(7906), 509–516.

67. Sjöström, P. J., and Häusser, M. (2006). A cooperative switch determines the sign of synaptic plasticity in distal dendrites of neocortical pyramidal neurons. Neuron, 51(2), 227–238.

68. Somogyi, P. (1977). A specific ‘axo-axonal’ interneuron in the visual cortex of the rat. Brain research, 136(2), 345–350.

69. Spratling, M. W. (2008). Predictive coding as a model of biased competition in visual attention. Vision research, 48(12), 1391–1408.

70. Stimberg, M., Brette, R., and Goodman, D. F. (2019). Brian 2, an intuitive and efficient neural simulator. elife, 8, e47314.

71. Tai, Y., Gallo, N. B., Wang, M., Yu, J. R., and Van Aelst, L. (2019). Axo-axonic innervation of neocortical pyramidal neurons by GABAergic chandelier cells requires AnkyrinG-associated L1CAM. Neuron, 102(2), 358–372.

72. Tasic, B., Yao, Z., Graybuck, L. T., Smith, K. A., Nguyen, T. N., Bertagnolli, D., … and Zeng, H. (2018). Shared and distinct transcriptomic cell types across neocortical areas. Nature, 563(7729), 72–78.

73. Tremblay, R., Lee, S., and Rudy, B. (2016). GABAergic interneurons in the neocortex: from cellular properties to circuits. Neuron, 91(2), 260–292.

74. Trubetskoy, V., Pardiñas, A. F., Qi, T., Panagiotaropoulou, G., Awasthi, S., Bigdeli, T. B., … and Lazzeroni, L. C. (2022). Mapping genomic loci implicates genes and synaptic biology in schizophrenia. Nature, 604(7906), 502–508.

75. Van de Cruys, S., Evers, K., Van der Hallen, R., Van Eylen, L., Boets, B., De-Wit, L., and Wagemans, J. (2014). Precise minds in uncertain worlds: predictive coding in autism. Psychological review, 121(4), 649.

76. Veres, J. M., Nagy, G. A., Vereczki, V. K., Andrási, T., and Hájos, N. (2014). Strategically positioned inhibitory synapses of axo-axonic cells potently control principal neuron spiking in the basolateral amygdala. Journal of Neuroscience, 34(49), 16194–16206.

77. Volk, D. W., Pierri, J. N., Fritschy, J. M., Auh, S., Sampson, A. R., and Lewis, D. A. (2002). Reciprocal alterations in pre-and postsynaptic inhibitory markers at chandelier cell inputs to pyramidal neurons in schizophrenia. Cerebral cortex, 12(10), 1063–1070.

78. Weiler, S., Guggiana Nilo, D., Bonhoeffer, T., Hübener, M., Rose, T., & Scheuss, V. (2023). Functional and structural features of L2/3 pyramidal cells continuously covary with pial depth in mouse visual cortex. Cerebral cortex, 33(7), 3715–3733.

79. Woodruff, A., Xu, Q., Anderson, S. A., and Yuste, R. (2009). Depolarizing effect of neocortical chandelier neurons. Frontiers in neural circuits, 3, 922.

80. Woodruff, A. R., McGarry, L. M., Vogels, T. P., Inan, M., Anderson, S. A., and Yuste, R. (2011). State-dependent function of neocortical chandelier cells. Journal of Neuroscience, 31(49), 17872–17886.

81. Wu, S. J., Sevier, E., Dwivedi, D., Saldi, G. A., Hairston, A., Yu, S., … & Fishell, G. (2023). Cortical somatostatin interneuron subtypes form cell-type-specific circuits. Neuron, 111(17), 2675–2692.

82. Xu, X., and Callaway, E. M. (2009). Laminar specificity of functional input to distinct types of inhibitory cortical neurons. Journal of Neuroscience, 29(1), 70–85.

83. Yizhar, O., Fenno, L. E., Prigge, M., Schneider, F., Davidson, T. J., O’shea, D. J., … and Deisseroth, K. (2011). Neocortical excitation/inhibition balance in information processing and social dysfunction. Nature, 477(7363), 171–178.

84. Yu, A.J. and Dayan, P. (2005). Uncertainty, neuromodulation, and attention. Neuron, 46(4), 681–692.

85. Zhang, C., Schneider-Mizell, C. M., Danskin, B. P., Swanstrom, R., Neace, E., Joyce, E., … & da Costa, N. M. (2026). Cell-type-specific parallel pathways in the canonical cortical microcircuit. bioRxiv.

