## Supplemental Figures for "Temporal Gating by Chandelier Cells Encodes Signed Prediction Errors"

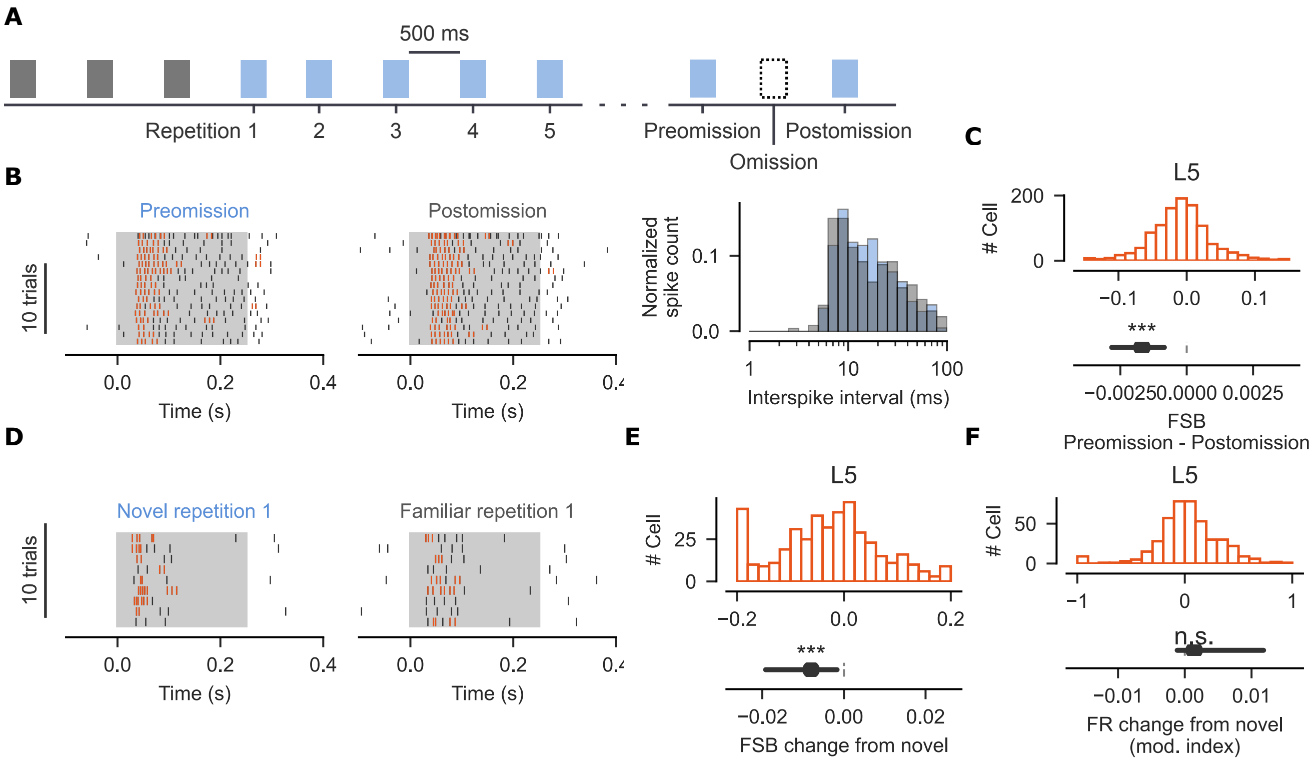


**Figure S1. L5 bursting scales proportionally to positive prediction error magnitude. Related to Figure 5.**

(A) Timeline of image presentations showing instances where some of the images are omitted from the sequence.

(B) Left: Raster showing the activity of the cell from Fig. 5A to presentation of the same image before omission (preomission) and directly after (postomission). Spikes that are part of a burst are shown in orange. The stimulus window is marked by grey background. Right: Histogram of interspike intervals during the stimulus presentation shown in Left. Spike count was normalised between the Pre-and Postomission conditions.

(C) Top: Histogram showing the change in FSB for individual cells from the Preomission to Postomission. Bottom: Confidence interval for the change. Mixed-effects model: $t_{(1074)} = -6.7$, $p < 10^{-10}$, CI = [-0.003, -0.001], n = 1075 cells from n = 98 recordings.

(D, E) As in (B) but comparing the response to the same stimulus when it was Novel (≤ 10 presentations) and Familiar (≥ 20 presentations). Mixed-effects model: $t_{(428)} = -3.6$, $p = 10^{-3}$, CI = [-0.019, -0.002], n = 429 cells from n = 49 recordings.

(F) Change in firing rate between Novel and Familiar image presentations. Firing rate was normalised to the activity during familiar presentations. Linear mixed-effects model: $t_{(428)} = 1.0$, p = 0.30, CI = [-0.001, 0.012], n = 429 cells from n = 49 recordings.

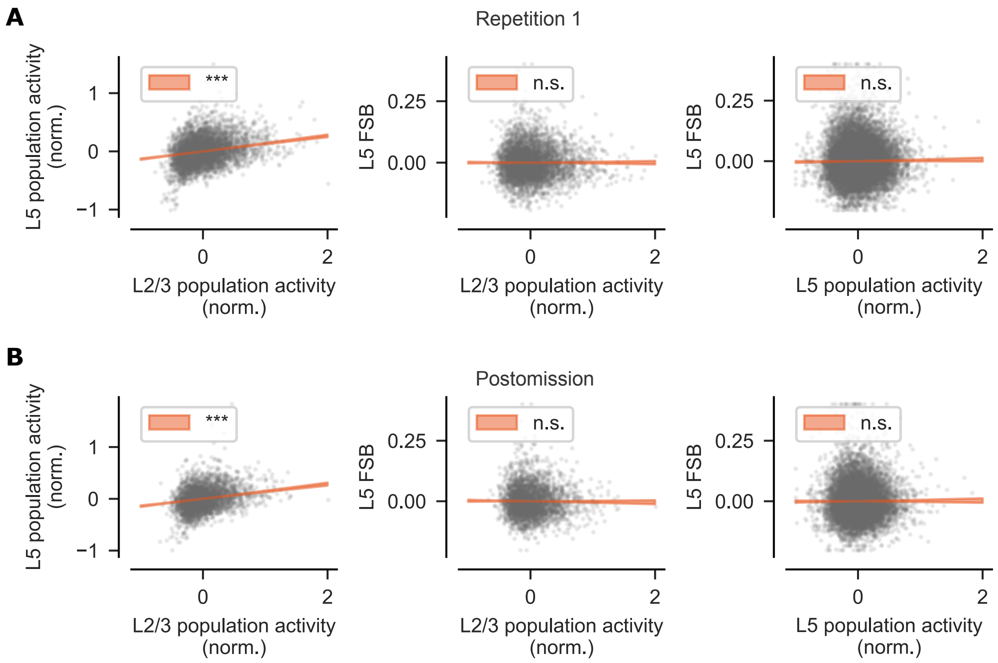


**Figure S2. L5 bursts are decoupled from L2/3 spiking. Related to Figure 5.**

(A) Left: Noise correlation of L2/3 and L5 population activity during the first image repetition. Linear mixed-effects model: $t_{(5221)} = 22.6$, $p < 10^{-112}$, CI = [0.12, 0.15], n = 5223 trials from n = 21 recordings. Middle: Noise correlation of L2/3 population activity and L5 FSB. Linear mixed-effects model: $t_{(4884)} = 0.0$, p = 0.99, CI = [-0.004, 0.004], n = 4886 trials from n = 21 recordings. Right: Noise correlation of L5 population activity and L5 FSB. Linear mixed-effects model: $t_{(19684)} = 1.6$, p = 0.11, CI = [-0.001, 0.007], n = 19686 samples from n = 86 recordings.

(B) As in (A) but calculated during the postomission stimulus. Left: linear mixed-effects model for L2/3 and L5 population activity noise correlation, $t_{(3743)} = 19.4$, $p < 10^{-83}$, CI = [0.13, 0.16], n = 3745 samples from n = 21 recordings. Middle: linear mixed-effects model for noise correlation between L2/3 population activity and L5 FSB, $t_{(3618)} = -0.95$, p = 0.34, CI = [-0.006, 0.002], n = 3620 samples from n = 21 recordings. Right: linear mixed-effects model for noise correlation between L5 population activity and L5 FSB, $t_{(14931)} = 0.6$, p = 0.55, CI = [-0.003, 0.006], n = 14933 samples from n = 86 recordings.
